# Mechanism of MEK1 phosphorylation by the N-terminal acidic motif-mediated asymmetric BRAF dimer

**DOI:** 10.1101/2025.09.26.678760

**Authors:** Yasushi Kondo, Judith Notbohm, Ignacio Navas Camacho, Gabriela Nagy-Davidescu, Thomas Mason, Jonas Mühle, Jörg Standfuss, Tina Perica

**Affiliations:** PSI Center for Life Sciences, Laboratory of Biomolecular Research, Paul Scherrer Institute, Villigen PSI, Switzerland; Department of Biochemistry, University of Zurich, Zurich, Switzerland; Biomolecular Mechanism and Structure Program, Life Science Graduate School Zurich, Zurich, Switzerland; Systems Biology Program, Life Science Graduate School Zurich, Zurich, Switzerland; Department of Chemistry, University of Zurich, Zurich, Switzerland

**Keywords:** MAPK/ERK signaling, BRAF, MEK, melanoma

## Abstract

The RAF/MEK/ERK signaling cascade regulates cell proliferation and differentiation and is frequently dysregulated in cancer. Approximately 90% of RAF-mutant cancers harbour mutations in B-type of Rapidly Accelerated Fibrosarcoma (BRAF). Its proto-oncogenicity has been attributed to a four-residue N-terminal acidic (NtA) motif. While a long-standing model proposes that the NtA promotes an activating asymmetric RAF dimerization, this idea has lacked structural support. Here, we present the first X-ray crystal structure of an NtA-mediated asymmetric BRAF dimer bound to its substrate Mitogen-activated Protein kinase Kinase (MEK1). Further cellular and biochemical analyses reveal that this asymmetric dimer is not an intermediate along the BRAF activation pathway, but rather a catalytically active state of the kinase. We validate this with the structure of BRAF bound to Ser222-phosphorylated MEK1, capturing the complex in a post-catalytic state. This combination of structural and cellular analysis establishes the relevance of NtA-driven asymmetry in BRAF activation and resolves a long-standing disconnect between RAF cancer genetics and structural biology.

## INTRODUCTION

Rapidly Accelerated Fibrosarcoma (RAF) Ser/Thr kinases initiate a mitogen-activated kinase cascade by doubly phosphorylating the activation loop of the mitogen-activated protein kinase kinase (MEK1) at positions Ser218 and Ser222,^1,2^ which in turn activates Extracellular Signal Activated Kinase (ERK). This RAF/MEK/ERK kinase cascade controls cell fate in metazoans by regulating proliferation, differentiation, and survival.^3^ Mutations that lead to uncontrolled signaling in this pathway are thought to drive tumorigenesis in ∼40% of human cancers.^4^ Activation of human RAFs is thus tightly regulated by a series of events, including phosphorylation/dephosphorylation, membrane localization, and conformational change.^5^ There are three human RAF paralogs, A-, B-, and CRAF with shared domain architecture (**Figure 1A**). In unstimulated cells, RAF and MEK kinases form stable cytoplasmic 1:1 heterodimers, where RAF is stabilized in an inactive conformation.^6,7^ Upon mitogen stimulation, GTP-bound small GTPase RAS (RAS•GTP) binds RAF and promotes RAF kinase domain dimerization that leads to an activated form of the kinase (**Figure S1**).^8,9^ An oncogenic variant of BRAF (Val600→Glu, BRAF-V600E) that is active independent of this multi-step mechanism involving RAS-binding followed by BRAF dimerization^10^ is responsible for approximately 50% of human melanomas.^11^

**Figure 1.**
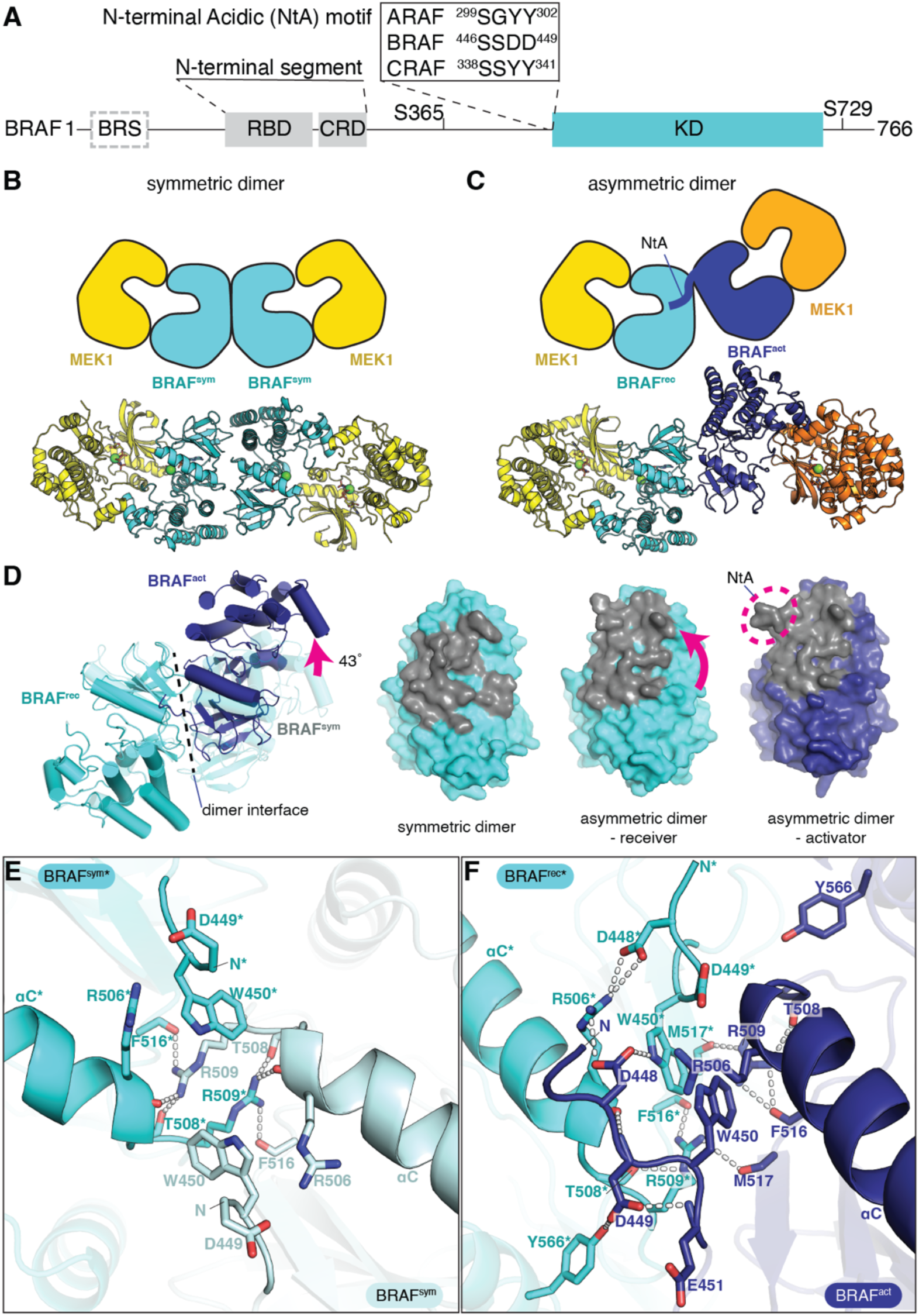
Two crystal forms of the BRAF:MEK1 heterotetramer reveal a symmetric and an asymmetric arrangement of the BRAF kinase domain dimer. (A) Domain architecture of BRAF. BRS – BRAF specific domain; RBD – Ras binding domain; CRD – cysteine-rich domain; NtA – N-terminal acidic motif; KD – kinase domain. (B) A symmetric and (C) an asymmetric BRAF:MEK1 complex in schematic (top) and cartoon (bottom) representation. The asymmetric complex comprises a BRAF activator subunit (BRAF^act^) with an NtA motif (dark blue) and a BRAF receiver subunit (BRAF^rec^) (cyan). ATP-analog bound to BRAF and MEK1 kinase active sites is shown in stick, and the metal ions are in sphere representation. (D) Shift of the BRAF:BRAF dimeric interface between symmetric and asymmetric dimers. The BRAF^act^ subunit is rotated by 43° compared to one of the BRAF subunits in the symmetric dimer (pink arrow). Interface residues (gray) are shifting between the symmetric and asymmetric dimer. (E and F) Rewiring of intermolecular contacts between (E) the symmetric and (F) the asymmetric BRAF dimer. Amino acids belonging to BRAF^rec^ (cyan) are annotated with an asterisk. Residues Asp448 and Asp449 that form intermolecular contacts in the asymmetric dimer are part of the NtA motif.

Current structural information on RAF kinases available to date supports a model where homo- or heterodimerization between different RAF paralogs imposes an active kinase domain conformation on each of the RAF subunits in a symmetric manner (**Figure S1**).^12^ In the cell, monomeric BRAF is stabilized in a catalytically-incompetent conformation that closely resembles the prototypical ‘CDK/Src-inactive kinase state’.^13–15^ As it is stabilized in an inactive conformation, RAF can remain bound to ATP and its substrate MEK1 without phosphorylating it.^6,16^ When two RAFs form a side-to-side dimer,^9^ the dimer interface formation enforces an active kinase conformation on both subunits^17^ and with MEK1 bound to each BRAF molecule a MEK1:BRAF:BRAF:MEK1 tetramer is formed.^18^ Although a high-resolution structure of this tetramer shows BRAF exhibiting the main hallmarks of an active kinase,^18^ the BRAF conformation still lacks some key elements necessary for catalysis: The ATP-analog available in the solution is not bound to the BRAF active site, and the MEK activation loop is sequestered from the BRAF active site. This raises the question of the extent to which this structure represents the true final active conformation of BRAF.

In addition, the BRAF:MEK1 complex structure does not capture the role of a four-residue N-terminal acidic (NtA) motif, a conserved but less understood motif in RAF (**Figure 1A**). Although the motif is nowhere near the active site,^8^ replacing it with four alanine residues (AAAA) significantly decreases RAF kinase activity.^19^ More specifically, the AAAA mutant of RAF failed to transactivate the other RAF kinase in the dimer, suggesting a role for an asymmetric arrangement in the RAF dimer.^20^ Furthermore, replacing the NtA motif of CRAF (SSYY) with the one from BRAF that has two phosphomimetic aspartates (SSDD) increases CRAF’s transforming activity.^21^ This explained why BRAF, whose kinase domain is otherwise >75% identical in sequence to ARAF and CRAF, is the one mutated in ∼90% of cancer patients with RAF mutations.^11^ However, despite its functional importance, the mechanism of how these four amino acids activate RAF in cells remained unknown.

Here we present structures of the BRAF:MEK1 heterotetramer with an asymmetric arrangement of the two BRAF kinase domains captured before and after MEK1 Ser222 phosphorylation. Importantly, this asymmetric dimer is mediated by the NtA motif. We show that replacing the NtA motif with four alanine residues (SSDD to AAAA in BRAF) abolishes *in vitro* kinase activity even though a complex with 14-3-3 is formed, as well as that it decreases MAPK pathway activity in cells in a dose-dependent manner. Taking together the structural features and our functional analyses, we propose that the BRAF asymmetric dimer is an essential conformational state of the fully active BRAF.

## RESULTS

### Two crystal forms of the BRAF:MEK1 complex reveal distinct dimerization interfaces

To obtain a structure of the MEK1:BRAF:BRAF:MEK1 complex that captures the phosphotransfer reaction state, we set up the crystallization experiment with the following modifications from a previous study.^18^ First, we did not use a MEK allosteric inhibitor, as MEK inhibitors act by sequestering the MEK activation loop (MEK-AL), including MEK1 residues to be phosphorylated, Ser281 and Ser222, from the BRAF active site.^22^ Second, we used a previously established BRAF kinase domain construct developed for bacterial expression,^23^ but with two modifications: 1) to preserve the native BRAF:MEK1 interface, we reverted the two mutations at the MEK1 interface (Phe667 and Tyr673) to wild type and 2) to overcome the inhibitory action of MEK1 and ATP on BRAF,^6,16^ we added the Val600Glu mutation to stabilize the active conformation of BRAF. The crystallization attempts yielded two unique crystal forms (**Figures 1B and 1C**): Crystal form I, obtained at neutral pH (6.5∼7.5) shows a symmetric side-to-side BRAF dimer, while the crystal form II, obtained at higher pH (8∼9) shows an asymmetric BRAF dimer (**Table S1**).

The BRAF:MEK1 structure obtained from crystal form I with the symmetric BRAF dimer is overall comparable to the previously reported structure by Haling et al.^18^ BRAF exhibits a conformation of an active kinase, which includes the inward shift of the αC helix and the extended activation loop, positing the catalytic residues in place (**Figure S2A**). However, unlike in the Haling structure, in our symmetric dimer the active sites in both BRAF subunits are occupied by a non-hydrolysable ATP-analogue AMPPNP (**Figure S2B**). At 2.9 Å resolution with anisotropic scaling,^24^ we could build AMPPNP and one metal ion at each of the two BRAF active sites (**Figure S2B**). The metal ion is associated with the Asp594 side chain of the ^594^DFG^596^ motif and β- and γ-phosphates of AMPPNP, which structurally corresponds to the high affinity M1 site in the prototypical Protein Kinase A (PKA) (**Figure S2C**).^25^

MEK1 in our complex with the symmetric BRAF dimer also shows differences from the Haling structure. The three-residue ^188^HRD^190^ motif is highly conserved among kinases and involved in the formation of the catalytic center.^26^ One of the MEK1 subunits in our structure has His188 flipped out, which creates a pocket filled by Phe209 (**Figures S3A and S3B**). This rewiring of the HRD motif residues is coupled to a shift in the MEK1 activation loop, as well as a 13° rotation of the BRAF:MEK1 interface compared to the Haling structure (**Figure S3C**).

The differences between our and Haling symmetric structures notwithstanding, in both structures the two MEK1 serine residues that are to be phosphorylated are at least 13 Å away from each of the BRAF active sites. In addition, the BRAF P-loop, a conserved kinase motif that takes part in the nucleotide and substrate binding and catalysis,^27^ remains disordered even with the nucleotide present, a notable difference from the structure of PKA,^28^ the canonical active kinase. In summary, although the ATP analog is present in the BRAF active site and the MEK inhibitor is absent, the conformation between BRAF and MEK1 in the complex with a symmetric BRAF dimer still does not appear to be fully compatible with the phosphotransfer reaction to Ser218 or Ser222 of MEK1.

Crystal form II, on the other hand, has two sets of BRAF:MEK1 heterodimers in the asymmetric unit, with an asymmetric BRAF dimer interface (**Figure 1C**). We further improved the resolution of this crystal form from 3.1 to 2.6 Å by using ^Δ36^MEK1 instead of ^Δ60^MEK1. ^Δ36^MEK1 contains a negative regulatory helix (residues 43-59),^29^ and both constructs were co-crystallized with BRAF in indistinguishable crystal forms and without significant differences between the structures (**Table S1**; **Figure S4**). In this asymmetric arrangement, the structures of the two BRAF subunits are similar to each other and to the BRAF molecules in the symmetric dimer, but the two BRAF kinase domains are rotated 43° around the dimeric interface with respect to the interface in the symmetric dimer (**Figure 1D**), revealing a surprising alternative arrangement of the BRAF dimer.

### The N-terminal acidic motif mediates the BRAF asymmetric dimer interface

In the asymmetric complex, the BRAF dimer interface maintains the apparent two-fold symmetry and involves largely the same residues as in the symmetric complex. However, the intermolecular residue-residue contacts are substantially reorganized between the two dimers. In the symmetric BRAF dimer, Arg509 is centrally positioned and forms an intermolecular handshake motif by interacting with the main chain carbonyl groups of Arg506 and Thr508 (**Figure 1E**).^30^ In contrast, within the asymmetric dimer, the Arg509 side chain folds back to intramolecularly bridge the main chain carbonyl groups of Thr508 and Phe516. Simultaneously, it forms an intermolecular hydrogen bond with the main chain carbonyl group of Met517 (**Figure 1F**). In the symmetric dimer, the highly conserved Trp450^20^ stacks against the Arg509 from the opposing subunit (**Figure 1E**). In the asymmetric dimer, the indole ring of Trp450 rotates towards the Trp450 from the other BRAF subunit, enabling a domain rotation that brings the two residues closer (**Figure 1F**). These structural changes highlight an unexpected plasticity of the BRAF dimeric interface.

Notably, our structure of the BRAF:MEK1 tetramer with an asymmetric BRAF dimer reveals that the aspartate residues of the NtA motif (^446^SSDD^449^) of BRAF form extensive dimeric interface contacts with residues in the conserved ^506^RKTR^509^ motif^31^ of the other BRAF subunit (**Figure 1F**). Specifically, the Asp448 side chain forms hydrogen bonds with the side chains of Arg506 and Trp450. The backbone amide group of Asp449 interacts with the carbonyl group of Arg506. These interactions are asymmetric: the NtA motif of the other BRAF in the dimer is exposed to the solvent (**Figure 1F**).

This asymmetric BRAF dimer arrangement is reminiscent of the model proposed by Hu et al. where one kinase in the dimer functions as an activator for the other, receiver kinase.^20^ According to this model, the NtA motif is essential for the activator function. Based on our structure, we propose that the BRAF monomer whose NtA motif forms extensive contacts at the dimer interface acts as the activator (BRAF^act^), while the other subunit functions as the receiver kinase (BRAF^rec^). In line with this nomenclature, we henceforth refer to the two BRAF subunits in the symmetric dimer as BRAF^sym^.

### The asymmetry in the BRAF active sites – catalytic BRAF^rec^ and nucleotide-free BRAF^act^

All three dimeric BRAF conformations: BRAF^sym^, BRAF^act^, and BRAF^rec^ exhibit many canonical features of an active kinase (**Figures 2A-2D**).^32^ However, in several conformational features, including interactions with the AMPPNP nucleotide and metal ions, the BRAF^rec^ subunit of the asymmetric dimer appears closer to the canonical active kinase (**Figure 2E**) than the BRAF subunits of the symmetric dimer (BRAF^sym^).

**Figure 2.**
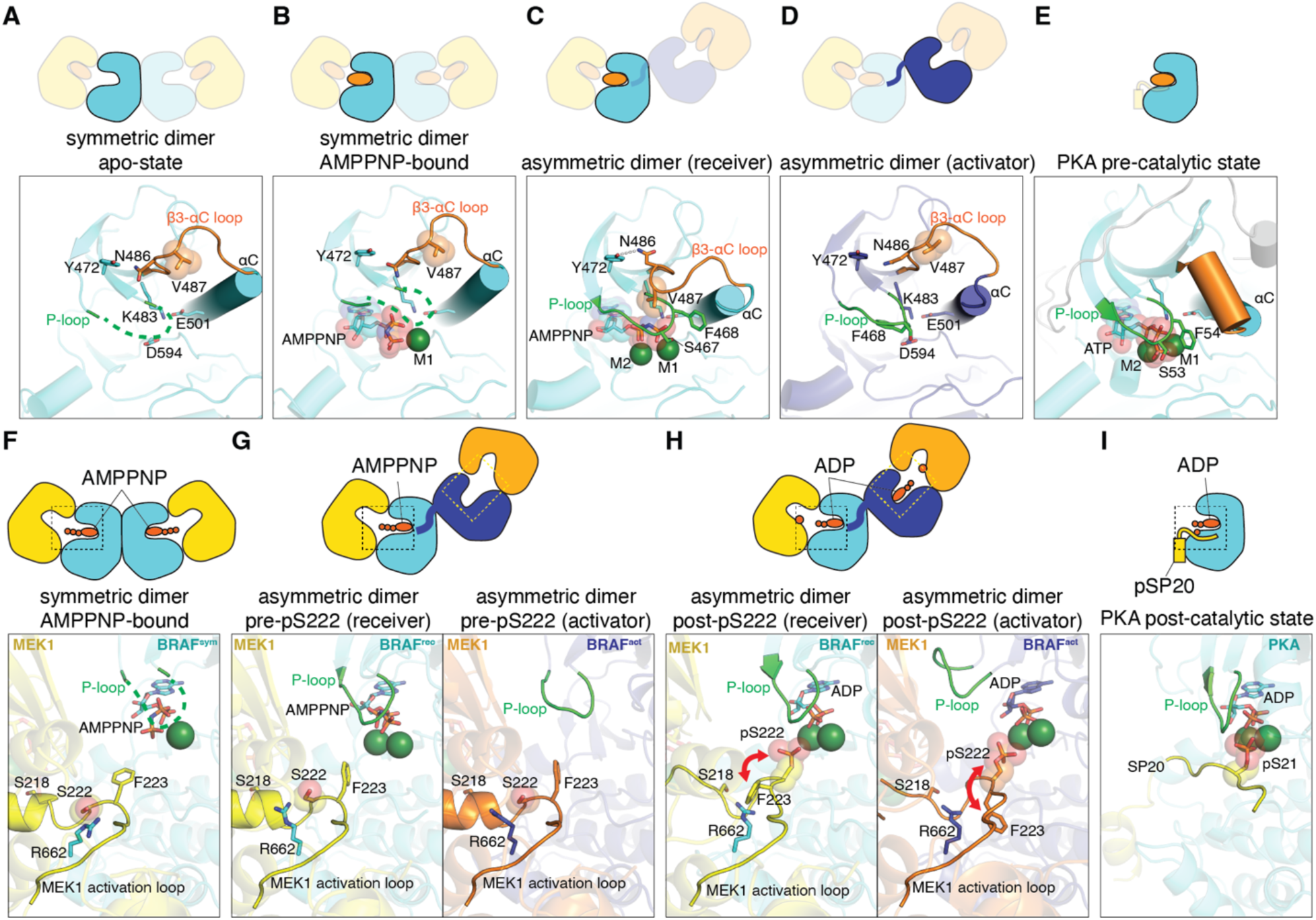
Asymmetry in the BRAF active sites. (A-D) X-ray crystal structures of active sites of BRAF kinase domains in different conformations. (A) The symmetric dimers in an apo-state (the Haling structure^18^) and (B) AMPPNP-bound state (this study) have their P-loops disordered (green dashed line). (C) In BRAF^rec^, the P-loop (green) interacts with the ATP-analog (AMPPNP) nucleotide. The β3-αC loop of BRAF^rec^ folds on top of the P-loop to stabilize it. (D) BRAF^act^ has an empty ATP-binding pocket. Conformational variability of the β3-αC loop is highlighted by the positions of Val487. (E) The PKA model (PDB ID: 1ATP) represents the canonical pre-catalytic state of a kinase.^28^ (F-H) Conformations of MEK1 activation loops. The MEK1 activation loop (MEK-AL, residues 208-230) and two phospho-acceptors (Ser218 and Ser222) are highlighted (yellow). The conformational change of the MEK-AL before and after Ser222 phosphorylation is highlighted by the movement of residues Ser222 and Phe223. (F) AMPPNP bound symmetric dimer, (G) AMPPNP bound asymmetric dimer, and (H) ADP-bound asymmetric dimer are shown. Red arrow indicates a position swap between Ser222 and Phe223. (I) PKA post-catalytic state co-crystallized with the product (pSP20) (PDB ID: 4IAF)^37^ is shown as a reference.

Striking features of the BRAF^rec^ active site are structural rearrangements in the β3-αC loop and the phosphate-binding P-loop (**Figure 2C**). Notably, the Val487 side chain in the β3-αC loop of BRAF^rec^ is shifted by 9 Å relative to the same residue in the symmetric dimer, stabilizing the closed P-loop. This shift is largely supported by the αC helix, which is positioned even more inward in BRAF^rec^ and BRAF^act^ than in BRAF^sym^ (**Figures 2B-2D and S5A**). This inward movement of the αC helix from opened to closed is a key step of kinase activation (**Figure S1**) and is likely promoted by the NtA-mediated dimeric interface interactions in the asymmetric dimer.

Furthermore, in the BRAF^rec^, AMPPNP is coordinated by two magnesium ions positioned by side chains of Asp594 and Asn581 (**Figure S5B**). The incorporation of the second metal ion at the M2 site in the asymmetric dimer thus appears to reflect a maturation of the BRAF catalytic site towards a fully active conformation. With AMPPNP bound and the P-loop closed, the BRAF^rec^ active site closely resembles that of active PKA (**Figures 2C, 2E, and S5B**),^28^ except that AMPPNP is positioned deeper within the active site of BRAF^rec^ (**Figure S5C**).

Unexpectedly, whereas both BRAF^sym^ and BRAF^rec^ contain an ATP-analog bound to their active sites (**Figures 2B and 2C**), the BRAF^act^ active site remains unoccupied (**Figure 2D**). The N-lobe of BRAF^act^ is rotated inward even further with respect to the BRAF^rec^ structure (**Figure S5D**). The β3-αC loop of the BRAF^act^ N-lobe adopts a conformation similar to the one in BRAF^sym^, leading to P-loop distortion that occludes the ATP-binding pocket (**Figure 2D**). The BRAF^act^ hinge loop displays a conformation different to one in any other BRAF structure. The Cys532 of the hinge loop and His585 of the β7-β8 loop switch positions, displacing the main chain amide group of Cys532, which typically forms a hydrogen bond with the N1 atom of ATP (**Figure S5E**).^33^ In contrast to the catalytically competent BRAF^rec^ active site stabilized by the BRAF^act^ NtA, the BRAF^act^ active site is constricted, supporting its non-catalytic role in the complex.

### Singly phosphorylated MEK1 in complex with BRAF trapped by crystallization

We hypothesize that the active site of BRAF^act^ remains unoccupied because it is too small to fit an ATP-analog. To test whether BRAF^act^ could fit the smaller ADP, we crystallized the complex in the presence of ADP. Indeed, the asymmetric BRAF:MEK1 complex accommodated ADP in both BRAF active sites (**Figure S6A**; **method details, Table S1**). The overall structure of the ADP-bound complex closely resembles that of the AMPPNP-bound structure with several key differences. The BRAF^act^ hinge loop reverts to the canonical conformation, and the P-loop adopts an open configuration, creating space for the nucleotide and forming contacts with the MEK1 N-lobe. This unusual P-loop conformation is likely driven by the N-lobe rotation seen in the AMPPNP-bound form (**Figure S5B**) and is further stabilized by ADP binding.

Surprisingly, we observed electron density at both MEK1 activation loops, into which we could model a phosphorylated serine (pSer222) (**Figure S6B**). This unexpected capture of the singly phosphorylated MEK1 in the crystal is likely due to trace ATP contamination in commercial ADP^34^ combined with crystallization in high concentration of Ca^2+^, which slows down nucleotide exchange.^35^ This interpretation is supported by an *in vitro* kinase assay showing MEK1 phosphorylation upon addition of ADP (**Figure S6C**). The resulting complex represents an intermediate state where MEK1 Ser222, but not Ser218, is phosphorylated. Trace levels of ATP were likely crucial for this serendipitous capture of the post-catalytic structure, as higher ATP concentration would lead to double (pSer218 pSer222) phosphorylation and thus dissociation of the BRAF:MEK1 complex.^18^

Upon Ser222 phosphorylation, the MEK-AL undergoes a dramatic conformational change. The originally α-helical segment becomes extended, and the Ser222 side chain rotates toward the BRAF active site, switching position with Phe223 of MEK1 (**Figures 2F-2H**). These changes are coupled to repositioning of BRAF Arg662, which switches from interactions with the Met219 carbonyl group to the Asp217 side chain and Ser218 carbonyl group (**Figures S7A-S7D**). Interestingly, molecular dynamics study of the BRAF:MEK1 symmetric dimer reported similar movements,^36^ suggesting that the observed structural transition reflects a conformational selection mechanism.

The distance of Ser222 to the β-phosphate of ADP in BRAF decreases from 13.9 Å in the unphosphorylated complex to 7.3 Å (BRAF^rec^) and 7.7 Å (BRAF^act^) after phosphorylation. These distances remain longer than the corresponding 6.2 Å in the PKA:SP20 post-catalytic complex (**Figure 2I**),^37^ reflecting differences in substrate peptide positioning (**Figure S7E**). Phosphorylation-induced rearrangement in the MEK-AL also propagates to the DFG motif at the N-terminal end of the activation loop. The metal-coordinating residue Asp208 of the DFG motif changes to a different rotamer and binds a second metal ion (**Figure S7F**). This transition of the DFG motif during unphosphorylated to phosphorylated state change corresponds to a shift from an inactive BLBplus to an active BLAminus conformation that has been observed across kinases.^38^

### BRAF N-terminal Acidic motif is essential for BRAF activity in cells

Our BRAF:MEK1 tetramer structure with an asymmetric BRAF arrangement supports the receiver-activator model, where the NtA-motif of the activator kinase forms dimeric interface contacts and, in turn, activates the other kinase in the dimer (the receiver).^20^ To assess the relevance of the receiver-activator model for BRAF function in cells, we overexpressed full length wildtype BRAF N-terminally tagged with mEGFP, as well as BRAF variants abrogating activator, receiver, or both roles at the same time, in human HEK293T cells. The activator role was abrogated by mutating the NtA motif to four alanines, resulting in a BRAF-AAAA variant that can only act as a receiver kinase. The receiver role was abrogated by an alanine to phenylalanine mutation (Ala481Phe) where the bulky side chain blocks ATP binding, making the BRAF-A481F variant catalytically incompetent and thus only capable of performing the activator role. Consequently, the double mutant, BRAF-AAAA-A481F acts neither as an activator nor as a receiver.^20^ We analyzed the effects of abrogating the activator or receiver functions individually and in combination on downstream pathway activity upon stimulation with human recombinant Epidermal Growth Factor (EGF) using flow cytometry (**Figures 3A and 3B**).^39,40^

**Figure 3.**
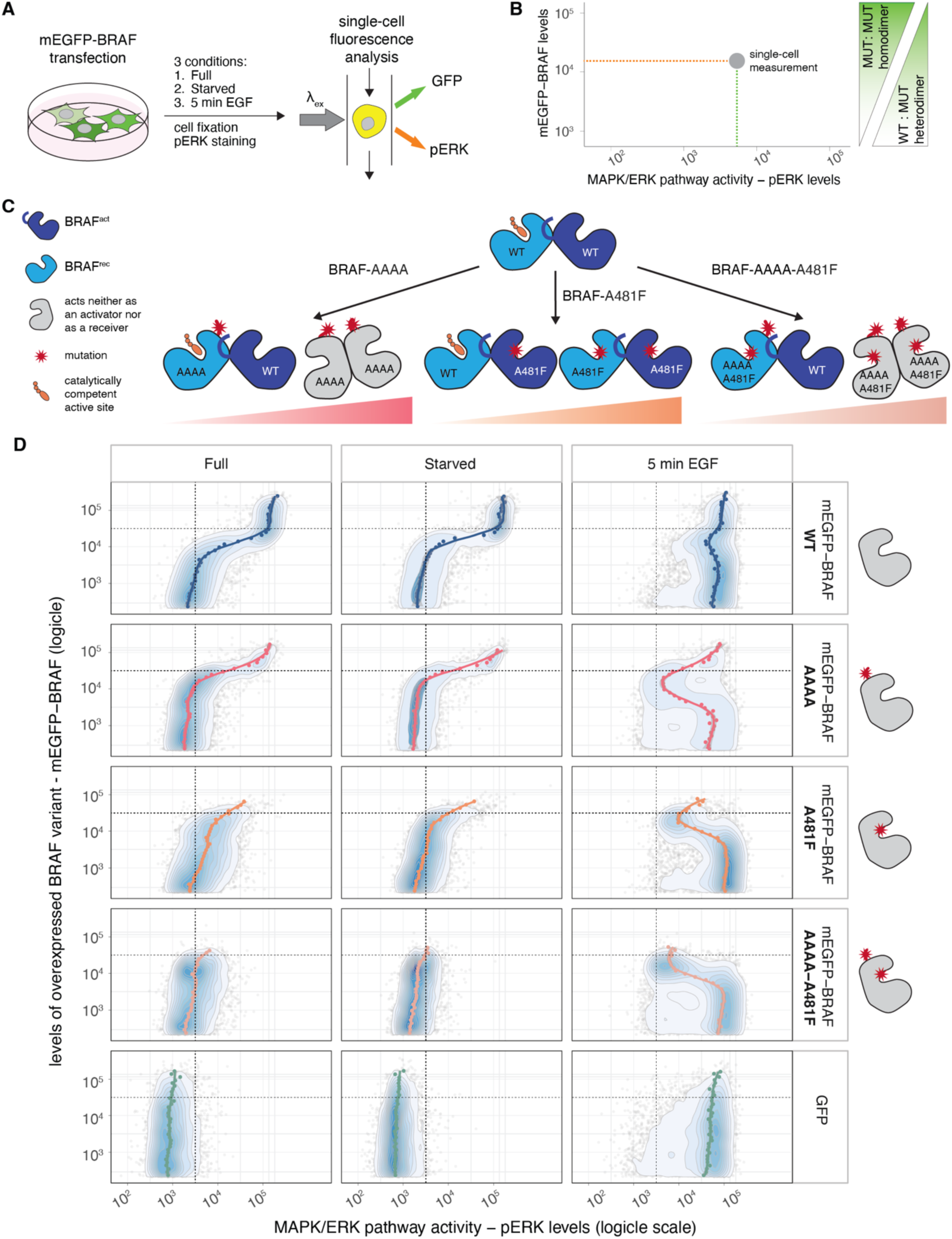
Dose response of MAPK/ERK activity (pERK levels) in HEK293T cells to activator and receiver-abrogating mutations in BRAF. (A) Experimental setup scheme. Flow cytometry analysis of single cells expressing different mEGFP-BRAF variants, analyzed under three conditions: Full - cells grown in media with 10% fetal bovine serum (FCS); Starved - cells starved for 2 hours in media without FCS; and 5 min EGF - cells stimulated with 30 ng/ml human recombinant Epidermal Growth Factor (EGF) for 5 minutes after 2 hours of starvation. After treatment, cells were stained for pERK, and levels of mEGFP- BRAF and pERK were measured for each cell by flow cytometry. (B) Higher expression levels of exogenous mEGFP-BRAF variants increase the total levels of RAF kinases in the cells, but also the ratio of MUT:WT heterodimers *versus* WT:WT endogenous RAF homodimers. Flow cytometry plots show how MAPK/ERK activity (pERK levels) respond to these different ratios for activator and receiver-breaking BRAF mutants. (C) In a cell expressing activator or receiver-abrogating BRAF mutants, three possible RAF dimers are formed: WT:WT, WT:MUT, or MUT:MUT. With increased mutant expression levels, the ratio of MUT:MUT dimers increases, and WT:WT decreases. Only WT:WT, WT:BRAF-AAAA, and WT:BRAF-A481F dimers have an active RAF kinase, highlighted by ATP in the kinase active site (orange). (D) Flow cytometry data collected as depicted in (A and B). Values are plotted on a logicle scale.^58^ Samples gated for single cells are shown as contour plots with outlier cells represented as individual points, where blue corresponds to highest density and white to lowest density of cells. The trend lines for each condition are colored by BRAF variant (blue for wild type, pink for BRAF-AAAA, orange for BRAF-A481F, and salmon for the double mutant, BRAF-AAAA-A481F) and represent the median pERK levels per each of the 50 GFP level bins of equal length. pERK activity is quantified with a pERK (pThr202/pTyr204) primary antibody followed by a secondary Alexa Fluor 647 conjugate antibody. Protein levels of overexpressed BRAF variants are quantified via fluorescence of the amino-terminal monomeric EGFP tag. The vertical and horizontal dashed lines are added to aid comparison between plots.

We used flow cytometry to leverage the diversity in the population of transiently transfected HEK293T cells. Namely, overexpression of BRAF variants in the background of endogenous RAFs results in a mix of WT:WT, WT:MUT and MUT:MUT dimers whose relative proportions are dependent on the levels of overexpressed BRAF variant in each individual cell (**Figure 3C**). In brief, we optimized transient expression of mEGFP-BRAF variants (**Figure S8**, **Table S2**) and then starved cells for 2 hours before stimulating them with EGF. We then fixed and stained the cells for pERK. In this way we were able to correlate pERK levels with BRAF expression levels in full media (DMEM + 10% FCS), starved (DMEM without serum), or stimulated conditions (5 min of 30 ng/ml EGF) and evaluate how abrogating activator and receiver roles affects MAPK/ERK signaling in a dose-dependent manner (**Figure 3**).

At low levels of exogenous mEGFP-BRAF, dimers of endogenous RAF dominate, and all variants behave similar to cells expressing GFP only, showing only slightly increased baseline pERK levels. At highest levels of exogenous mEGFP-BRAF, the cells overexpressing wild type BRAF show high pERK levels, even in serum-free starvation conditions. This population is smaller for variants abrogating the activator and receiver functions and absent for the double mutant variant in all three independent transfection replicates (**Figures 3D**, **S9, and S10**; starved condition). At the same time, the mutation abrogating only the receiver role (BRAF-A481F) leads to higher pERK baseline levels compared to the wild type already at low expression levels (**Figure 3D**; full condition), reminiscent of paradoxical activation^41–43^ and signaling by BRAF class 3 oncogenic mutants.^44^ The double mutant (BRAF-AAAA-A481F), however leads neither to cells with high pERK levels in starved condition at high expression, nor to paradoxical activation at low expression. Taken together, we conclude that the additive effect of the AAAA-A481F double mutant confirms the importance of the asymmetric arrangement of BRAF dimers (**Figure 3C**).

Upon EGF stimulation, we observe the most striking differences between cells expressing wildtype or mutant BRAF at levels that are just below those at which pERK is constitutively phosphorylated. At these levels, a large proportion of cells expressing either single (AAAA or A481F) or double (AAAA-A481F) BRAF mutant fail to activate pERK (**Figure 3D**; 5 min EGF condition). We observed the same effect when we quantified pMEK instead of pERK (**Figure S11**) and we could corroborate this by imaging pERK-stained cells, which showed that high exogenous BRAF expression is mutually exclusive with high pERK levels for mutant variants but not for wildtype BRAF (**Figure S12**). We reasoned that the distinct inactive population of cells appears at exogenous variant expression levels at which inactive MUT:MUT dimers saturate the endogenous pool of RAS•GTP (**Figure S13A**).

At the same expression levels where we observe the phenotype in stimulated cells for the three BRAF mutant variants, we also observed a reproducible dip in pERK and pMEK levels in cells expressing wildtype BRAF (**Figure 3D**; 5 min EGF condition; **Figure S11**). As the dip was independent of the tag (**Figure S8B**) and was not observed in cells expressing GFP only, we postulate it is due to combinatorial inhibition^45,46^ where at high BRAF expression levels, formation of KRAS:BRAF and BRAF:MEK complexes inhibits formation of the KRAS:BRAF:MEK complex (**Figure S13C**). When we co-expressed exogeneous mCherry-KRAS with the mEGFP-BRAF, we observed neither the pERK dip nor the large population of cells that fail to activate pERK at high BRAF-AAAA levels (**Figures S13D and S14A**). However, we could still observe a dose-dependent effect of BRAF levels on pERK levels when we overexpressed BRAF-AAAA versus BRAF-WT (**Figure S14B**).

Our single cell resolution flow cytometry results agree with the bulk Western blot experiments based on which Hu et al.^20^ originally proposed the activator-receiver model. Their experiments were predominantly performed with truncated BRAF constructs without the KRAS-binding domain, where the measured downstream ERK activity was independent of upstream signals and pERK levels were only quantified in the full media condition. When we quantified pERK levels in bulk with Western blot using our full-length constructs and a range of conditions (**Figure S15**), we only observed differences in full media conditions when we co-expressed KRAS but could observe clear decrease in pERK levels for the receiver, activator, and double mutant upon EGF stimulation (**Figure S15**).

In summary, the NtA motif is necessary for KRAS-mediated BRAF activation and removing NtA has a dose-dependent effect on signaling, as BRAF without the NtA can still act as a receiver, but not as an activator kinase. In line with that, a variant that can act neither as an activator nor as a receiver (BRAF-AAAA-A481F) has an additive effect. Our results support the importance of the BRAF asymmetric dimer formation in MAPK/ERK signaling in the cell.

### The NtA motif mediated asymmetric dimer is formed by active BRAF on the membrane

We next investigated the role of the NtA-mediated asymmetric dimer in RAF activation in more detail. Namely, the asymmetric dimer could represent an intermediate state along the activation path of BRAF, which spans from the autoinhibited monomeric form to the KRAS-bound, fully active dimeric form (**Figure S1**). Or it could represent either an alternative or the final active form of the BRAF kinase, as suggested by the structure of the receiver molecule (**Figure 2C**). To place the asymmetric dimer along the BRAF activation path, we tested how NtA ablation in the BRAF-AAAA variant affects two key RAF activation steps: KRAS-mediated membrane localization and SHOC2:MRAS:PP1C-mediated pSer365 dephosphorylation.

To test if the asymmetric dimer formation is necessary for KRAS•GTP-mediated membrane localization, we transfected HEK293T cells with mCherry-KRAS-IRES-eGFP-BRAF constructs, encoding either wild type or Gly12Asp variants of KRAS, and wild type, AAAA or Arg188Leu variants of BRAF. A construct with an Internal Ribosome Entry Site (IRES) ensured that each cell has sufficient levels of KRAS to localize all GFP-BRAF to the plasma membrane. As expected, we observed that the wild type BRAF, unlike the Arg188Leu variant that cannot bind KRAS, localizes to the plasma membrane in a KRAS-G12D-dependent manner (**Figures 4A and 4B**). Importantly, ablation of the NtA motif (BRAF-AAAA) did not appear to affect the KRAS-G12D mediated localization of BRAF. This is in agreement with the studies on CRAF, whose NtA needs to first be phosphorylated to become acidic, as CRAF NtA phosphorylation has been shown to occur at the plasma membrane.^19,47^ Taken together, we conclude that an NtA motif-mediated asymmetric dimer must be formed after RAFs associate via RAS•GTP to the membrane.

**Figure 4.**
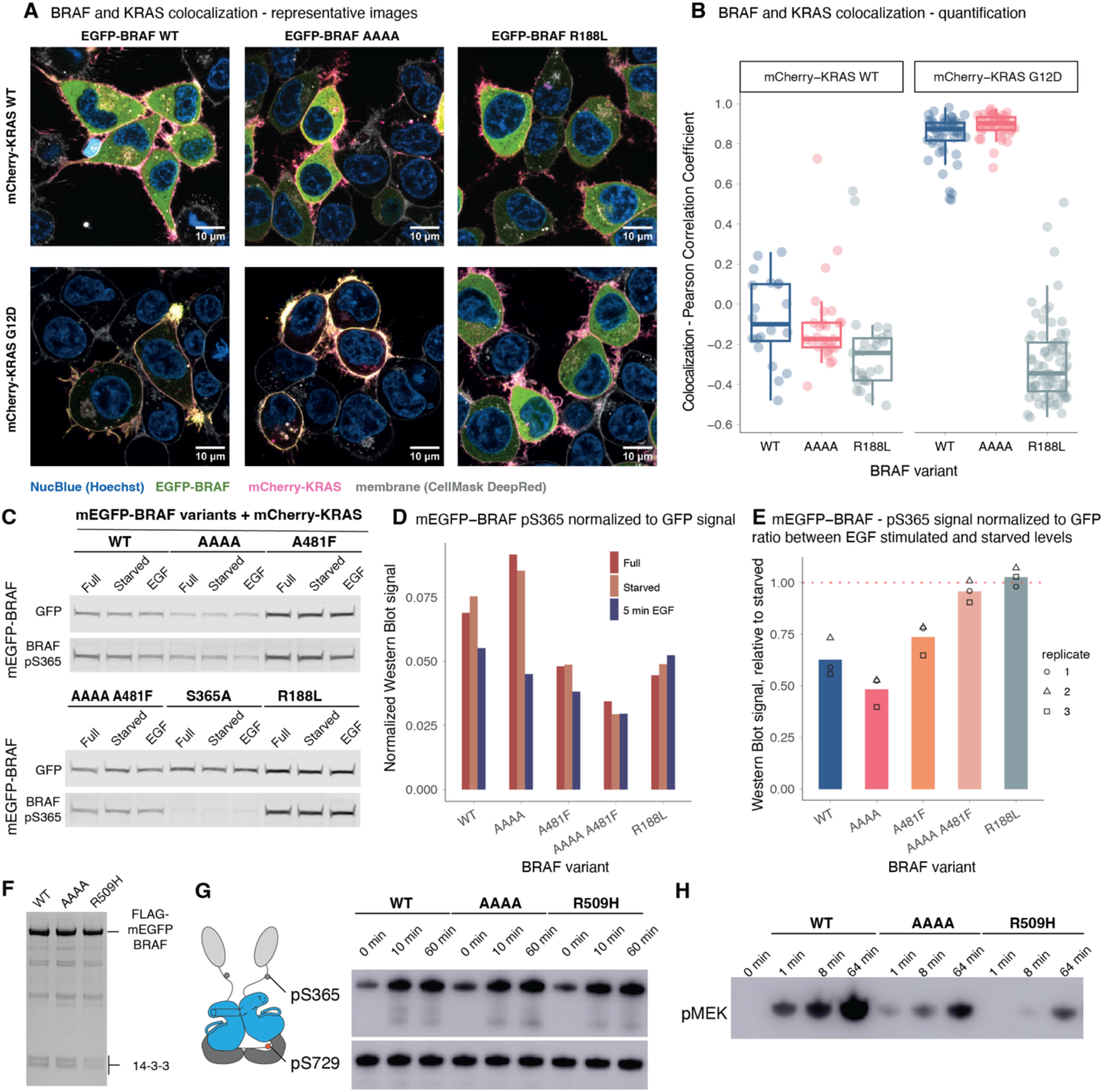
The N-terminal Acidic Motif (NtA) is not necessary for KRAS•GTP-mediated BRAF membrane localization nor for the activating BRAF pSer365 dephosphorylation. (A) Representative confocal microscopy images of HEK293T cells transfected with a mCherry-KRAS-IRES-EGFP-BRAF plasmid expressing different variants of BRAF and KRAS. BRAF is localized to the plasma membrane in a KRAS•GTP (KRAS-G12D) dependent-manner. Colocalization of EGFP-BRAF (green) and mCherry-KRAS (pink) on the membrane is indicated by bright yellow. (B) Quantification of colocalization (n = 20 to 70 cells) measured as Pearson correlation between BRAF and KRAS within a membrane-cytoplasm region of interest. (C) Example Western blot (replicate 2, same as in Figure S13C) quantifying levels of pSer365 in overexpressed mEGFP-BRAF variants co-expressed with mCherry-KRAS. Levels of mEGFP-BRAF variants and BRAF pSer365 were quantified with an anti-GFP and CRAF pSer259 antibody that can detect overexpressed pSer365 BRAF, respectively. Samples were collected for HEK293T cells grown in medium with 10% FCS (Full), in cells starved for 2 hours in media without FCS (Starved), and in cells stimulated after 2 hours starvation with 10 ng/ml of EGF for 5 minutes (EGF). GFP and pSer365 levels were detected from the same respective bands enabling quantitative comparisons. (D) Quantification of pSer365 signal normalized to GFP signal from the blot shown in (C). (E) Ratio of pSer365/GFP levels between starved and stimulated (5 min EGF) conditions. The bars show a mean of three independent biological replicates (independent transfections) of Western blots shown in Figure S13C. Values from independent replicates are shown as points. (F) SDS-PAGE of purified FLAG-mEGFP-BRAF proteins. BRAF proteins expressed in HEK293T cells were purified using an anti-FLAG antibody. 10 pmol of each protein, as estimated by GFP absorption, was applied to the gel. (G) BRAF phosphorylation status analysis by susceptibility to PKA. Purified FLAG-mEGFP-BRAF proteins were incubated with PKA and ATP. N- and C-terminal 14-3-3 binding interface phosphorylation sites (pSer365/pSer729) were monitored using phospho-specific antibodies. (H) Enzymatic activities of purified FLAG-mEGFP-BRAF variants measured by an *in vitro* kinase assay. Purified BRAF proteins were mixed with MEK1 and ATP, and MEK1 phosphorylation was detected by an anti-phospho-MEK antibody.

One of the final steps of BRAF (and CRAF) activation is presumably the dephosphorylation of pSer365 (pSer259 in CRAF) by the SHOC2:MRAS:PP1C (SMP) phosphatase complex.^48^ Knocking down the SHOC2 component of the phosphatase complex results in a low but sustained MAPK/ERK activity.^49^ This step follows the membrane localization via KRAS•GTP and is associated with the rearrangement of the interaction between RAF and 14-3-3, and RAF dimerization (**Figure S1**).

To determine whether the NtA motif, and in turn the asymmetric dimerization is necessary for pSer365 dephosphorylation, we co-expressed mCherry-KRAS and different mEGFP-BRAF variants in HEK293T cells and quantified pSer365 phosphorylation upon stimulation with EGF. In line with previous work,^49^ we observed that EGF stimulation results in ∼30-40% decrease in Ser365 phosphorylation in overexpressed wild type BRAF, but not in the Arg188Leu variant that affects BRAF binding to KRAS•GTP, and thus membrane localization (**Figures 4C-4E and S16**). Interestingly, we also did not observe a decrease in Ser365 phosphorylation in the BRAF-AAAA A481F double mutant, but this variant also consistently showed lower pSer365 levels in the starved condition. Importantly, the BRAF-AAAA variant showed comparable decrease in Ser365 phosphorylation to the wild type BRAF. Therefore, we conclude that the NtA motif, and in turn the asymmetric dimer formation, are not necessary for removal of this inhibitory phosphorylation that is coupled to RAF dimerization and 14-3-3 interface rearrangement.

Our membrane localization and the Ser365 dephosphorylation data demonstrate that the activator-receiver asymmetric interaction is likely not an intermediate step along the RAF activation pathway, but rather an integral state of activated RAF on the membrane. If that is the case, we expect that removing the NtA would prevent BRAF from achieving its full enzymatic activity.

To test the enzymatic activity and the oligomeric state of the NtA mutant BRAF-AAAA, we overexpressed FLAG-mEGFP-BRAF wild type, AAAA, and Arg509His variants in HEK293T cells grown in full medium supplemented with 10% serum and purified the resulting BRAF-containing complexes. Under these conditions, previously shown to produce the BRAF symmetric dimer bound to the 14-3-3 dimer^7^ we observed that all the variants, wildtype BRAF, the NtA mutant BRAF-AAAA and the dimer interface mutant BRAF-R509H co-purified with 14-3-3 (**Figure 4F**).

We also interrogated the phosphorylation status of the BRAF:14-3-3 binding sites by detecting availability of free serine residues for PKA. The analysis revealed that the C-terminal Ser729 site was fully phosphorylated, while the N-terminal Ser365 was only partially phosphorylated and could, thus, in all the tested variants be further phosphorylated by addition of PKA (**Figure 4G**). As expected, we observed that purified wild type full length BRAF can phosphorylate MEK1 *in vitro*. Furthermore, the kinase activity is dramatically decreased by a dimeric interface Arg509His mutation. Consistent with earlier studies,^19^ the purified BRAF-AAAA dimer had detectable enzymatic activity that was, however, strikingly lower than that of the wild type BRAF (**Figure 4H**), indicating that the ability to form an asymmetric dimer via the NtA motif is an essential step in achieving full kinase activity.

Taken together, our functional assay data suggest that BRAF needs to assume an NtA motif-mediated asymmetric dimer conformation to catalyze the phosphotransfer reaction. Why the activated BRAF kinase domain dimer assumes two distinct conformations, the symmetric and the asymmetric one, awaits further investigations.

## DISCUSSION

Our work provides molecular-level insight into the sequence of events by which BRAF phosphorylates MEK1 (**Figure 5**). Based on our data, we propose a model where ATP is bound to the active site of a symmetric BRAF dimer, after which an asymmetric dimer is formed, accompanied by rearrangements of β3-αC and P-loop of BRAF and the MEK-AL. Only then can MEK1 Ser222 phosphorylation occur, and we could capture a post-pS222 BRAF:MEK1 conformation. As the asymmetric dimer co-crystallized with ADP has both MEK1 subunits phosphorylated, the roles of receiver and activator within the BRAF dimer are likely interchangeable.

**Figure 5.**
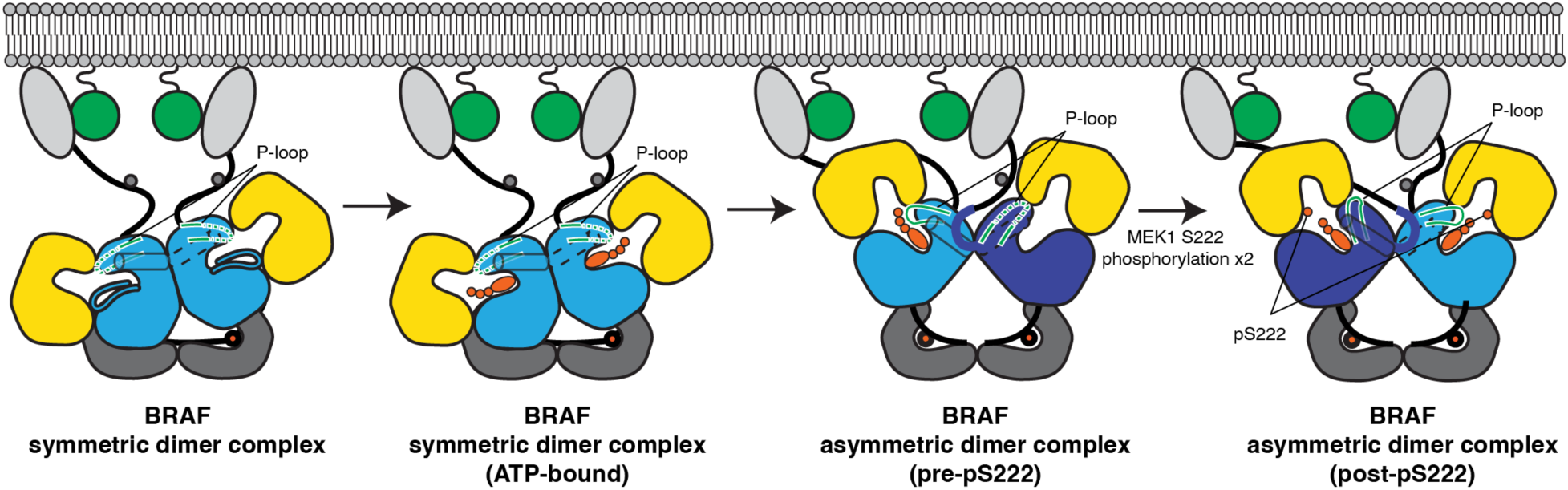
Schematic representation of the BRAF:MEK1 complex conformational change upon MEK1-Ser222 phosphorylation. For simplicity, structural elements of MEK1 (nucleotide, activation loop, αC helix, and P-loop) and the BRAF activation loop in the ATP-bound symmetric dimer and asymmetric dimer complexes are not shown. Color scheme is the same as in Figure S1.

An alternative interpretation of our data is that symmetric and asymmetric dimers represent distinct active states with higher and lower activity, respectively. However, the fact that the symmetric dimer can be crystallized in ligand-free state even when nucleotide is present in solution^18^ and that its P-loop remains disordered even in the AMPPNP-bound state suggests that the “open” active site of the symmetric dimer favors nucleotide release. Furthermore, an efficient nucleotide exchange is often a rate-limiting step in the kinase reaction.^50^ In the case of PKA, the N-lobe opens widely in the nucleotide-free state,^51^ but such a movement is sterically hindered in BRAF due to its dimer interface. We thus propose that wildtype BRAF cycles between asymmetric dimers for catalysis and symmetric dimers for nucleotide exchange.

Both Ser218 and Ser222 of the MEK-AL need to be phosphorylated for MEK1 to be activated and dissociate from the complex.^18,52^ Our structure captured the intermediate, singly phosphorylated MEK1 which showed that phosphorylated Ser222 is too bulky to fit back into the BRAF C-lobe pocket it occupies in the BRAF^sym^ conformation (**Figures 2F, 2G, S7B, and S7C**). The pSer222 rather remains close to the BRAF active site, where it would interfere with exchange of ADP for new ATP by clashing with its γ-phosphate (**Figure 2H and S7D**). We therefore expect the binding of the new ATP to trigger a conformational shift that might move Ser218, the next MEK1 residue to be phosphorylated closer to the BRAF active site. A similar mechanism has been proposed for arylalkylamine N-acetyltransferase, where product release is promoted by binding of the next acetyl-CoA molecule.^53^ Importantly, we cannot exclude that the BRAF:MEK1 complex dissociates between the Ser222 and Ser218 phosphorylation steps. Double phosphorylation is a conserved activation mechanism in MAPK pathways,^54^ and our model allows for either sequential^55^ or processive^56^ phosphorylation.

Our structures also reveal a progression in metal-binding modes within BRAF and MEK1 active sites. In the inactive, monomeric state, BRAF binds MEK1, an ATP analog and one metal ion, and adopts a CDK/Src-like^13–15^ or DFG-in BLBplus^38^ inactive conformation.^6,16^ Upon symmetric dimer formation, regardless of nucleotide or metal binding, BRAF transitions to an active, DFG-in and BLAminus conformation.^18^ When AMPPNP binds to the symmetric dimer, one metal ion occupies the M1 site, but the P-loop remains disordered. Finally, in the asymmetric dimer, the active site accommodates two metal ions, which we propose marks the transition to full catalytic competency. Interestingly, we observe an analogous structural propagation in MEK1: the unphosphorylated MEK1 adopts a BLBplus conformation with a nucleotide and one metal ion, while Ser222 phosphorylation induces a DFG shift to the BLAminus conformation, now coordinating two metal ions (**Figure S7F**). This model highlights the importance of phosphorylation-coupled structural rearrangements in MEK1 activation.

In cells, the symmetric BRAF dimer forms a stable complex with 14-3-3 via the C-terminal phosphopeptide (**Figure S1**).^6,7,12,16,57^ In this work we show that the asymmetric interaction between two BRAF kinase domains mediated by the NtA motif (^446^SSDD^449^) is compatible with the constitutive complexation with 14-3-3 (**Figures 4F-4H**). A transition from a symmetric to an asymmetric BRAF dimer requires a rotation that moves the C-terminal Leu721 residues of the two BRAF kinase domains from 43 to 66 Å apart (distances between Leu721 Cα atoms) (**Figure 1D**). The structural variability of the linker between the kinase domain and the 14-3-3-binding element of BRAF observed in multiple cryo-EM and X-ray crystallography structures would allow for such rearrangement.

Given their complex multi-step activation and regulation mechanisms, interpreting effects of overexpressing specific variants of signal transduction kinases in cells remains a challenge. Here we show that single cell approaches such as flow cytometry that can simultaneously quantify expression and phosphorylation levels of multiple signal transduction components can provide clearer insights than bulk measurements.

Finally, our work uncovers the NtA-mediated asymmetrical transactivation of RAF kinases. Importantly, the asymmetric transactivation model is the key for explaining the paradoxical activation of RAF by ATP-competitive BRAF inhibitors used in cancer treatment.^41–43^ The insights into understanding of RAF transactivation offered by our work might provide a new avenue for improving BRAF inhibitors for cancer therapy.

## RESOURCE AVAILABILITY

### Lead contact

Requests for further information and resources should be directed to and will be fulfilled by the lead contact, Tina Perica.

### Materials availability

Plasmids generated in this study are available upon request from the lead contact.

### Data and code availability

- Coordinates and structure factors have been deposited in the Protein Data Bank database under accession code 9RTQ (BRAF:MEK1 heterotetramer with the symmetric dimer interface with AMPPNP), 9RTR (BRAF:MEK1 heterotetramer with the asymmetric dimer interface with AMPPNP), and 9RTS (BRAF:MEK1 heterotetramer with the asymmetric dimer interface with ADP).
- All other data and analysis code are available at https://github.com/tinaperica/AsymmetricBRAF or as Supplementary Materials that are a part of this manuscript.

## SUPPLEMENTAL INFORMATION

**Figure S1.**
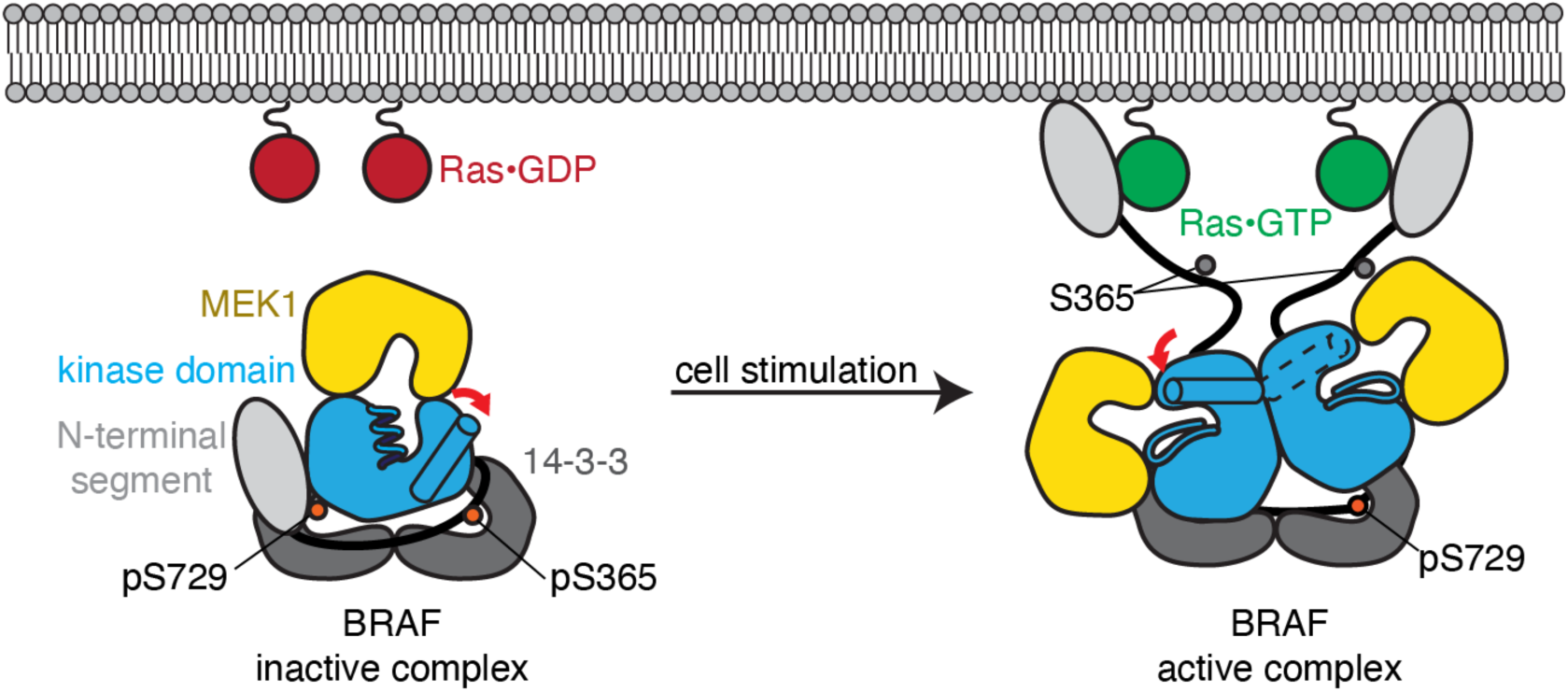
Multi-step activation of BRAF on the membrane. Current model of BRAF activation in the cell. An inactive BRAF complex comprises a BRAF monomer bound to MEK and a 14-3-3 dimer. In inactive, monomeric BRAF, the αC helix of the kinase domain is positioned outward and the activation loop is folded. Upon interaction with Ras•GTP on the membrane, the complex matures to an active form. The BRAF kinase domain forms a dimer and takes up an active conformation, where the αC helix is positioned inward and the activation loop is extended.

**Figure S2.**
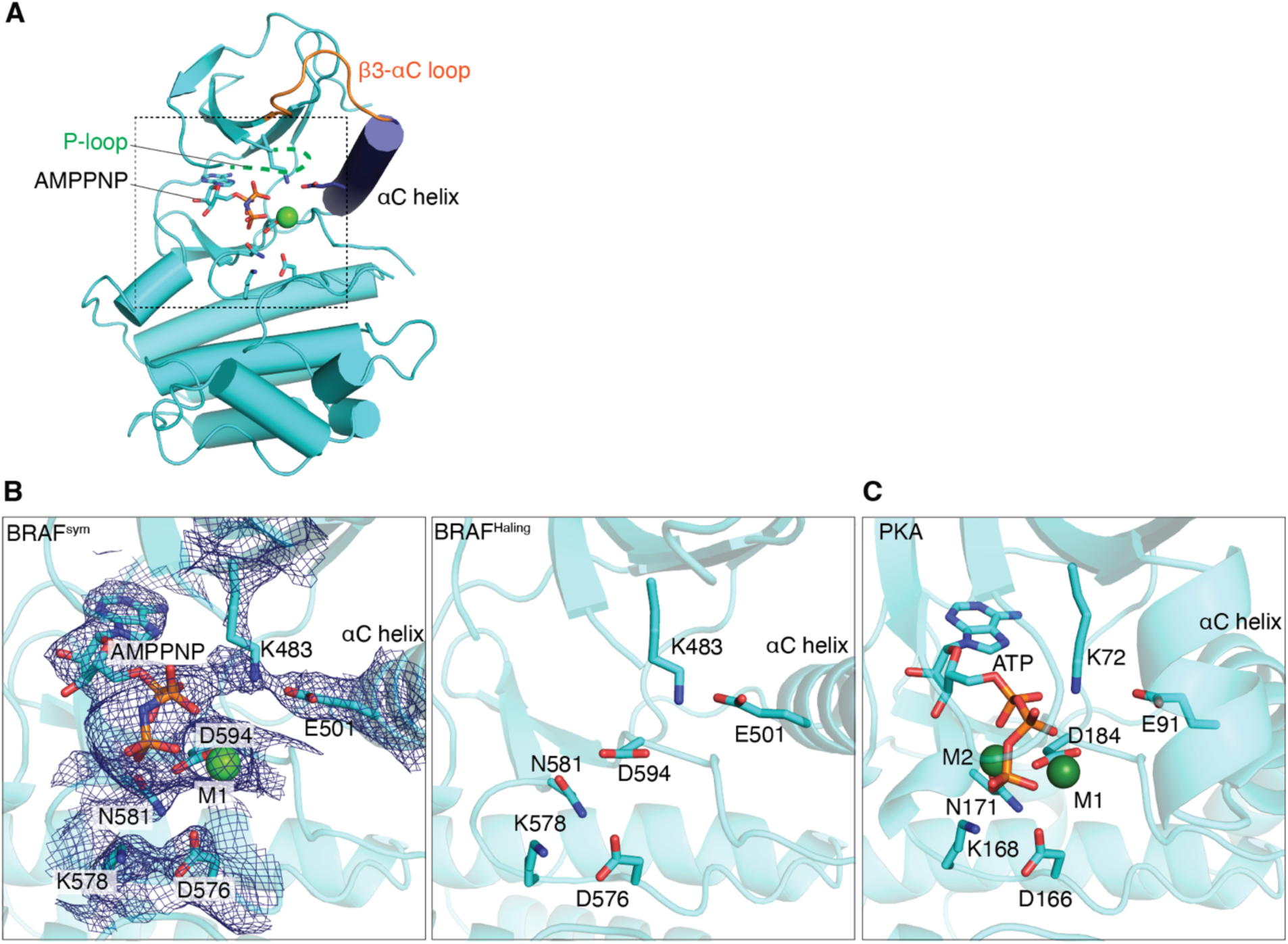
The active sites of the BRAF symmetric dimer. (A) The overall structure of the BRAF kinase domain. A single subunit of the AMPPNP-bound BRAF symmetric dimer exhibiting a canonical active kinase conformation. Key structural elements are highlighted: the β3-αC loop (orange); the P-loop (green); the αC helix (deepblue). A dashed box indicates the active site of the kinase. (B) The BRAF symmetric dimer structures with (left; this study) or without (right; Haling et al.^18^) ATP-analog show nearly identical conformations. The key amino acid residues in the active kinases are shown in stick. Based on crystallization conditions we modelled a calcium ion as seen in PKA,^68^ but in cellular conditions, we expect a magnesium ion at this position. The 2Fo-Fc map of the BRAF^sym^ is contoured at 1.0 sigma and carved around the AMPPNP and the key amino acid residues of the BRAF active site. (C) ATP and metal coordination in PKA (PDB ID: 1ATP). Amino acid side chains coordinating phosphate groups of ATP (Lys72 and Lys168) and metal ions (Asn171 and Asp184), as well as highly conserved active site residues Glu91 and Asp166 are shown in stick.

**Figure S3.**
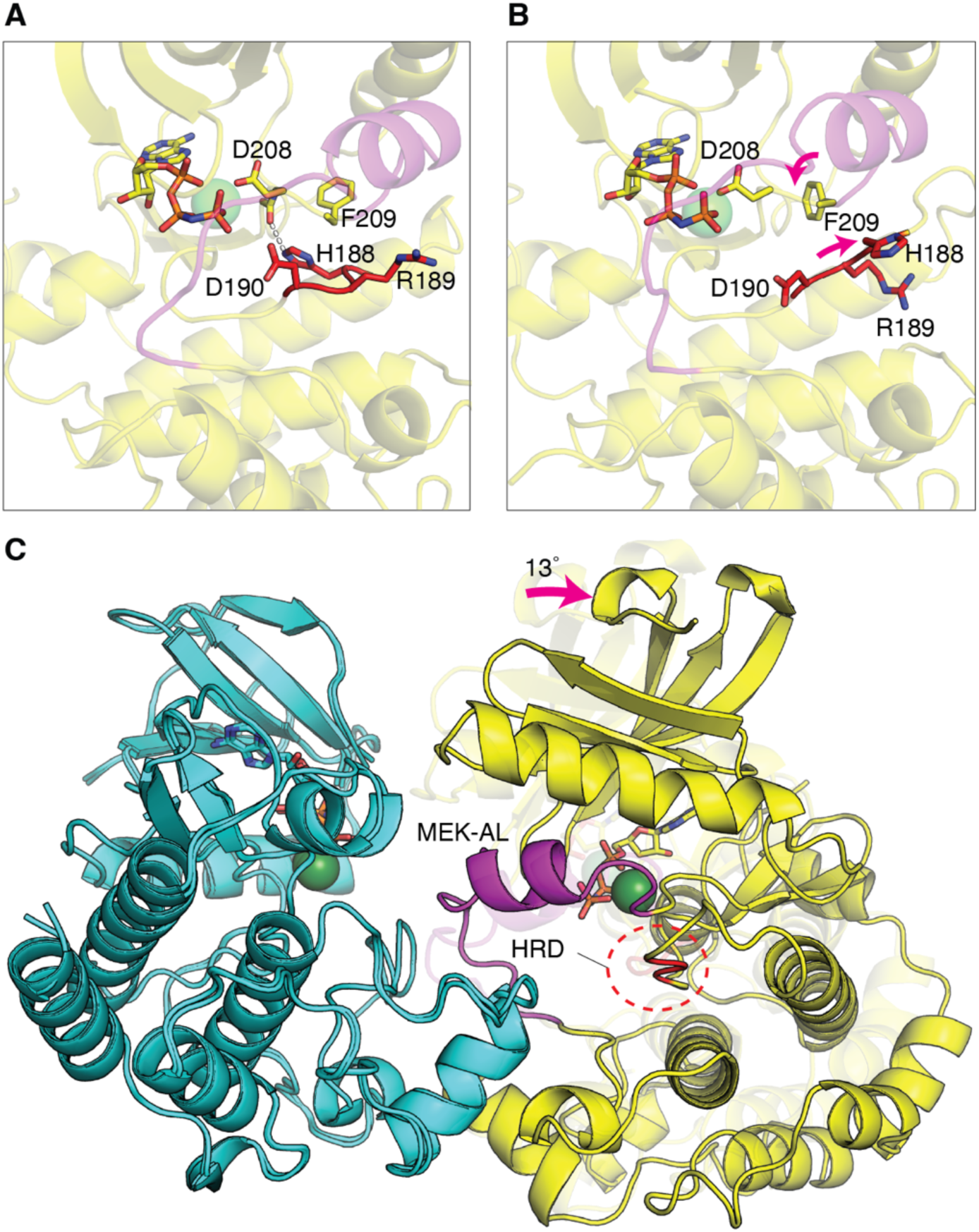
Alternative conformations of the MEK1 HRD motif in the BRAF:MEK1 structure with the symmetric BRAF dimer. (A and B) Two distinct conformations of the MEK1 ^188^HRD^190^ motif in the two MEK1 molecules bound to the symmetric BRAF dimer from this study. The three HRD residues are in red. Asp208 and Phe209 of the invariant DFG-motif are represented in stick. The MEK1 activation loop (MEK-AL) is in magenta. (A) One MEK1 subunit in the BRAF:MEK1 heterotetramer with the symmetric BRAF dimer shows a canonical conformation of the HRD motif, with the His188 sidechain interacting with the Cys207 mainchain carbonyl group. (B) The other MEK1 subunit in the heterotetramer shows an unusual HRD motif conformation. His188 flips out, and the Phe209 sidechain folds into the pocket created by this His188 flip. (**C**) Superposition of the BRAF C-lobes showing the MEK1 rotation in the symmetric dimer.

**Figure S4.**
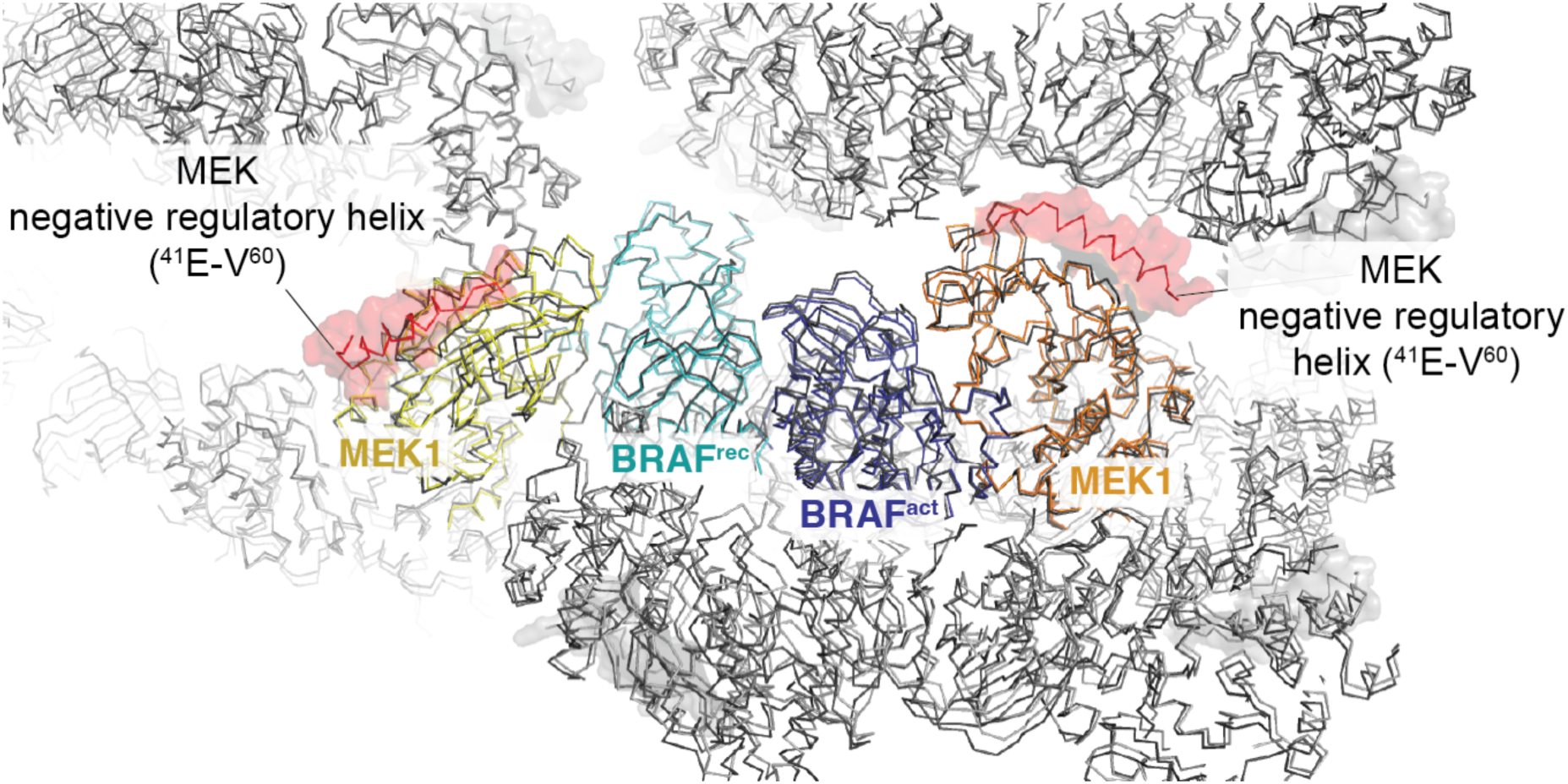
Structural comparison of the BRAF:MEK1 asymmetric dimers obtained by using two different MEK1 constructs, ^Δ36^MEK1 or ^Δ60^MEK1. The BRAF:MEK1 heterotetramer with ^Δ36^MEK1 is colored as in Figure 1C (yellow and orange), with the N-terminal helix additionally highlighted in red. The crystal structure with ^Δ60^MEK1 is shown in gray. The crystallographic symmetry related molecules are shown in light gray (^Δ36^MEK1) and dark gray (^Δ60^MEK1).

**Figure S5.**
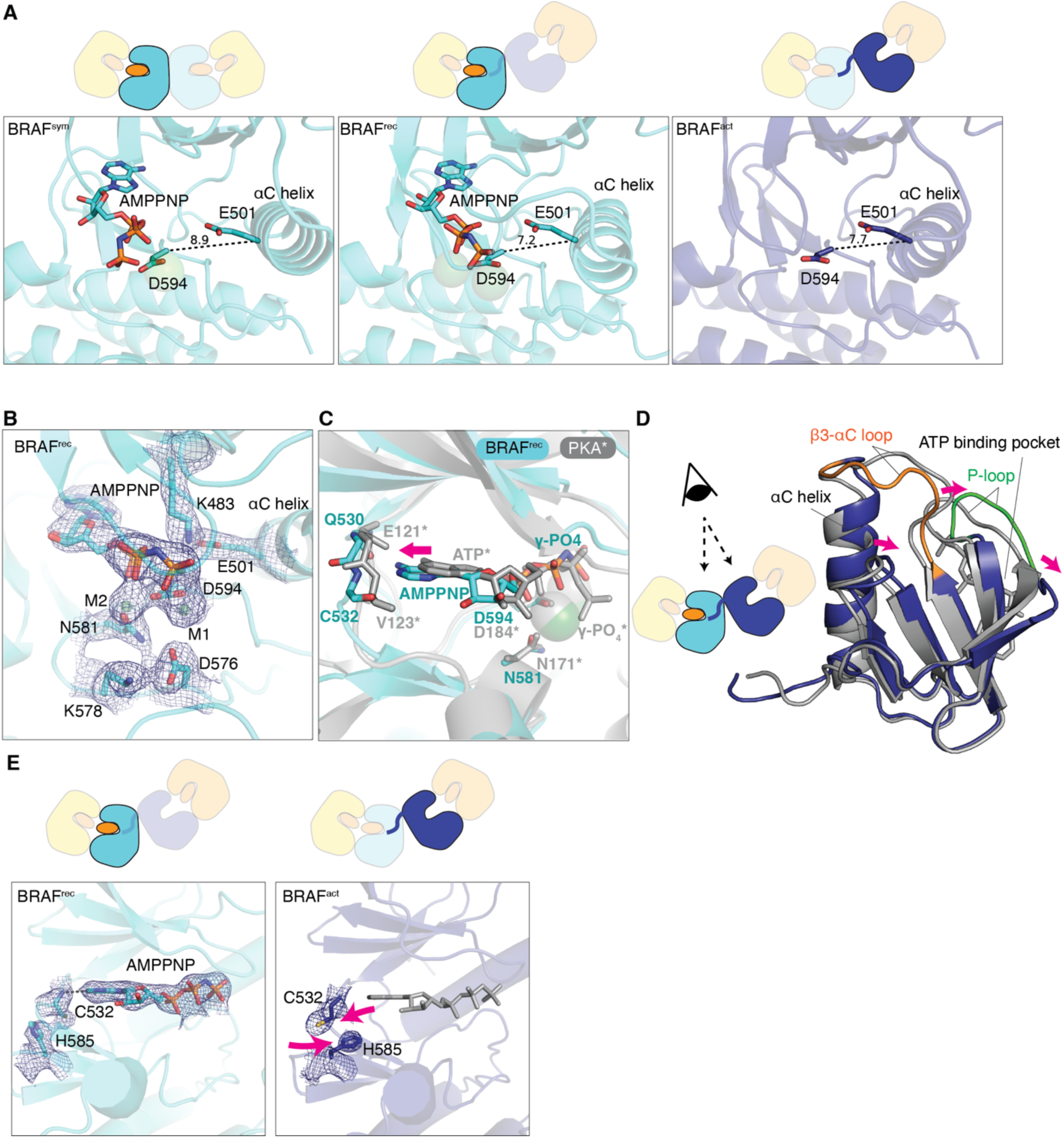
Structural comparison of BRAF^rec^ and BRAF^act^ subunits in the AMPPNP-bound asymmetric BRAF dimer. (A) The position of the αC helix in the three different BRAF subunit types, BRAF^sym^, BRAF^rec^, and BRAF^act^. The inward movements of the αC helices in the asymmetric dimer are highlighted by shorter distances between the Cα atoms of Glu501 (αC helix) and Asp594 (DFG motif): 8.9 Å in BRAF^sym^, compared to 7.7 Å in BRAF^act^ and 7.2 Å in BRAF^rec^ (B) The BRAF^rec^ active site with AMPPNP. The M1 metal primarily coordinates the γ-phosphate, while M2 bridges the β- and γ-phosphates. The 2Fo-Fc map of the BRAF^sym^ is contoured at 1.0 sigma and carved around the AMPPNP and the key amino acid residues in the BRAF active site. (C) Nucleotide positions in BRAF^rec^ and PKA. AMPPNP-bound BRAF^rec^ and pre-catalytic PKA structures were superposed using the C-lobe region (BRAF residues 532-723 and PKA residues 122-293). The nucleotide and the hinge loop of BRAF^rec^ are shifted away from the substrate binding site compared to PKA (pink arrow). Dashed lines highlight matching atoms represented in stick. (D) Top view of BRAF^rec^ and BRAF^act^ N-lobes. The two BRAF subunits making the asymmetric BRAF dimer are superposed via their C-lobes (residues 532-723). BRAF^act^ is in dark blue and BRAF^rec^ in gray. (E) Constricted ATP-binding pocket in the BRAF^act^ subunit of the asymmetric BRAF dimer. Two panels compare BRAF^rec^ and BRAF^act^ ATP-binding pockets. Gray dashed line indicates the hydrogen bond formed between the adenine ring and the hinge loop. AMPPNP molecule of BRAF^rec^ is superposed to BRAF^act^ and shown in gray. The 2Fo-Fc maps of the BRAF^rec^ and BRAF^act^ are contoured at 1.0 sigma and carved around the AMPPNP and the key residues in the BRAF^rec^ active site.

**Figure S6.**
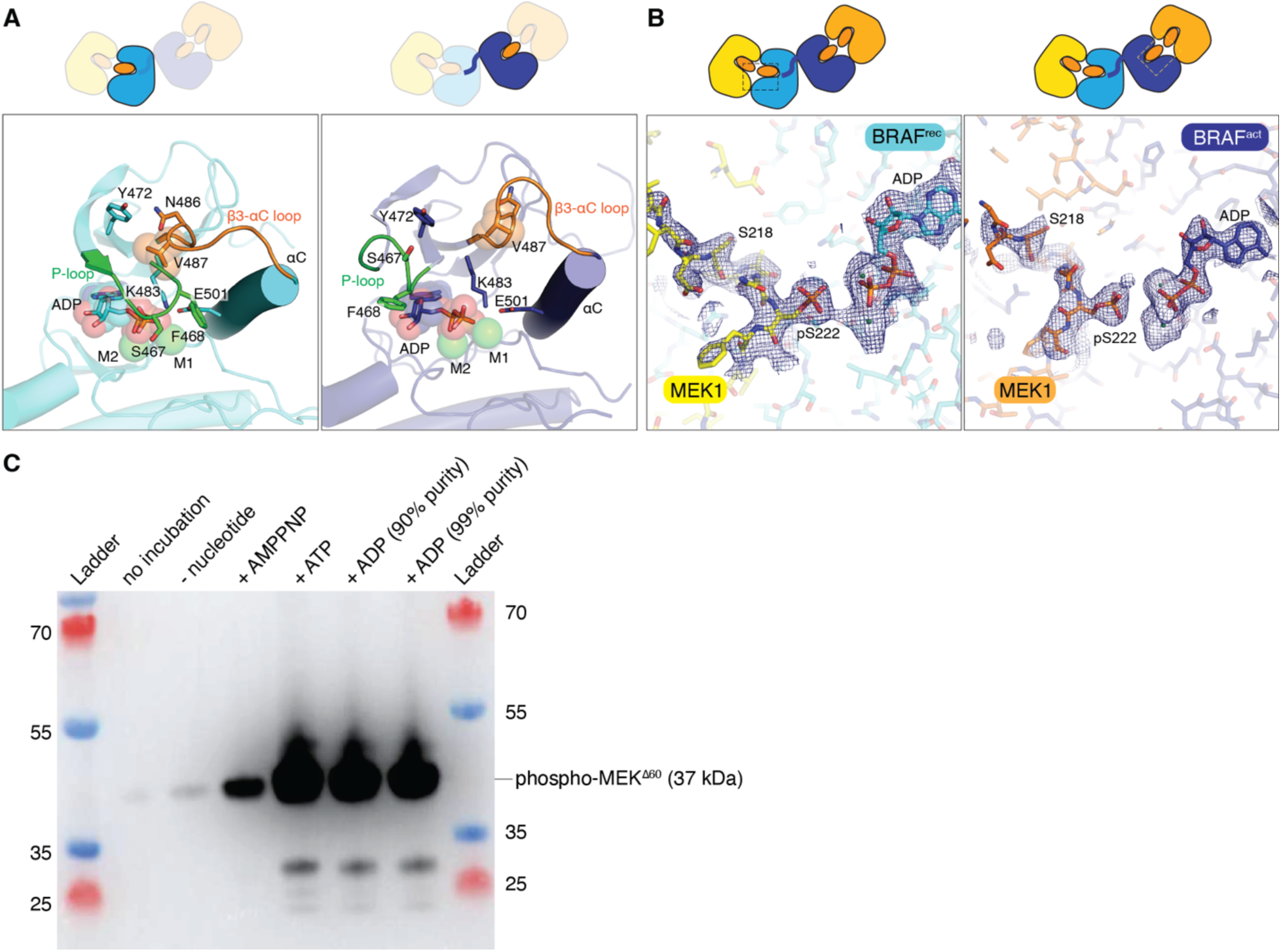
Structure of the BRAF:MEK1 asymmetric dimer co-crystallized with ADP. (A) ADP molecule bound to the BRAF^rec^ and BRAF^act^ active sites. Color schemes are the same as in Figure 2 (BRAF^rec^ (cyan) and BRAF^act^ (dark blue)). (B) Electron density map of the MEK1 activation loop (MEK-AL) including the Ser222 phosphorylation site. The 2Fo-Fc map is contoured at 1.0 sigma and carved around the MEK-AL and the ADP bound to the BRAF active site. (C) *In vitro* kinase assay results using different commercially-available ADP nucleotides. ^Δ60^MEK1 was incubated with the purified BRAF kinase domain and the different nucleotide reagents. Phospho-MEK1 was detected with an anti-phospho-MEK antibody. Due to trace contaminations with ATP, efficient MEK1 phosphorylation can be observed even when using commercial ADP with 99% purity.

**Figure S7.**
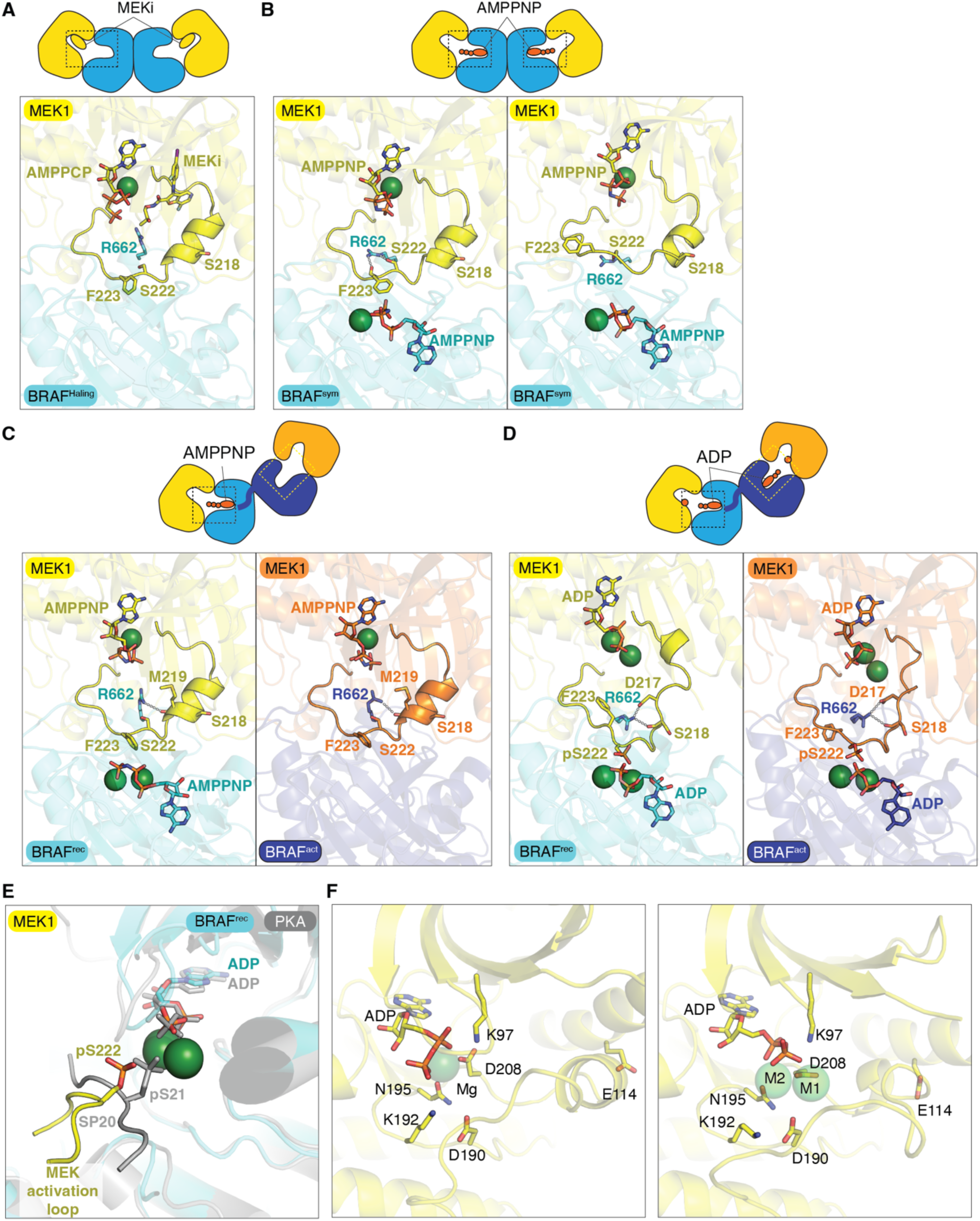
Conformational changes in the MEK1 activation loop upon Ser222 phosphorylation. BRAF:MEK1 crystal structures of the (A) symmetric dimer with the MEK inhibitor, G-573 (Haling structure; PDB ID: 4MNE),^18^ (B) the symmetric dimer with AMPPNP (this study, PDB ID: 9RTQ), (C) the asymmetric dimer with AMPPNP (this study, PDB ID: 9RTR), and (D) the asymmetric dimer with ADP and pSer222 (this study, PDB ID: 9RTS). Metal ions bound to the nucleotides are shown as green spheres. Hydrogen bonds are highlighted with dashed lines. The nucleotides bound to the MEK1 kinase domains are omitted from the schematic for simplicity. (E) The comparison of substrate peptide positions between PKA and BRAF. The BRAF asymmetric dimer with pSer222-MEK1 and the post-catalytic structure of the PKA:SP20 complex^37^ are superposed via their C-lobes. The PKA and the PKA substrate, SP20, are shown in gray. (F and G) Structural changes in the MEK1 active site. (F) An unphosphorylated MEK1 kinase domain bound to Mg•ADP (PDB ID: 3EQD)^29^ has a Src/CDK-like or BLBplus inactive conformation. (G) The singly phosphorylated MEK1 in the BRAF:MEK1 asymmetric dimer has a BLAminus active conformation.

**Figure S8.**
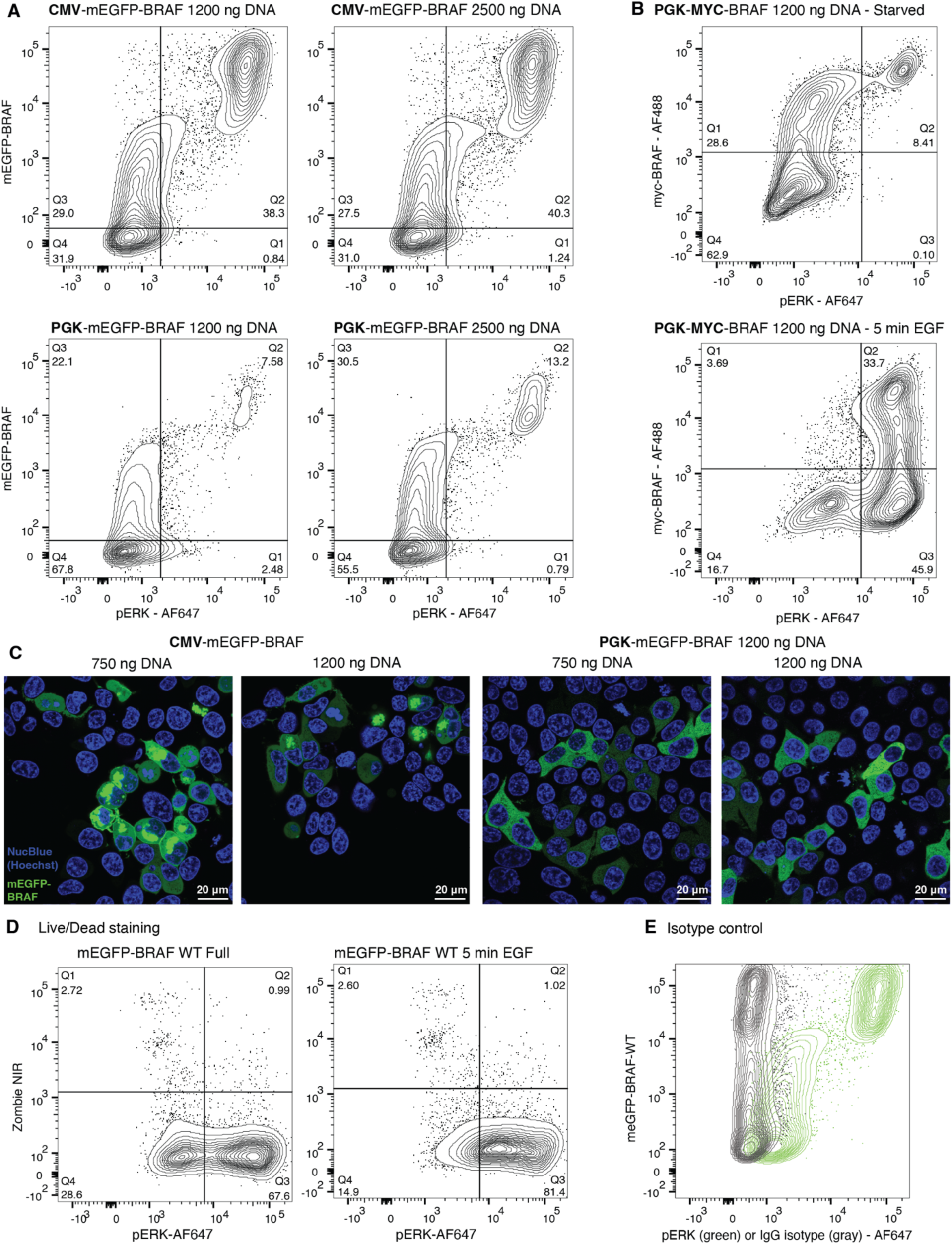
Under many conditions, BRAF overexpression results in a population of cells with constitutively high pERK levels, regardless of serum-free starvation. (A) Flow cytometry analysis of single cells expressing mEGFP-BRAF from a strong (CMV) or a weak (PGK) promoter^69^ and transfected with two different amounts of DNA, stained for pERK. Data shows the effect of promoter and transfection conditions on BRAF expression levels and on pERK levels in starve conditions. Samples gated for single cells are shown as contour plots with outlier cells represented as individual points. Gates shown were set on a non-transfected control (vertical) and a non-transfected starve control (horizontal). When mEGFP-BRAF is expressed from the CMV promoter, a substantial proportion of cells (quadrant Q2, ∼40%) have high pERK levels even after 2 hours of serum-free starvation. This problem is not alleviated by transfecting with less DNA (1200 ng instead of 2500 ng) but it is by using a weaker PGK promoter. pERK is quantified with a pERK (pThr202/pTyr204) primary antibody followed by a secondary Alexa Fluor 647 conjugate secondary antibody. Protein levels of overexpressed BRAF variants are quantified via fluorescence of the amino-terminal monomeric mEGFP tag. (B) Flow cytometry analysis of single cells expressing BRAF-WT with mEGFP exchanged for a myc-tag, in starved and 5 min EGF conditions. Gates shown were set on non-transfected controls (vertical) and a non-transfected starve control (horizontal). When the mEGFP tag is replaced with a smaller c-myc-tag, high BRAF expressing cells still show high pERK levels in serum-free conditions, indicating that it is the overexpression of BRAF rather than the mEGFP tag that causes this non-physiological response. (C) Live cell confocal imaging of HEK293T cells overexpressing mEGFP-BRAF from two different promoters, CMV or PGK, after transient transfection with different amounts of DNA. Transient overexpression of mEGFP-BRAF from the CMV promoter in HEK293T leads to spherical cells with observable GFP clusters in the cytoplasm. Nuclear DNA is stained with NucBlue (Hoechst, blue) while mEGFP-BRAF is in green. (D) Anti-pERK antibody does not specifically stain dead cells and cell viability is comparable between the full and stimulated conditions. Gates used in the plot were set on a Live/Dead control sample (with 50% of cells killed prior to staining) for Zombie NIR, and the starve sample for pERK-AF647. ∼3% of cells are dead after starvation and stimulation, comparable to the full media sample. (E) Isotype control to rule out unspecific binding of the pERK antibody, especially in cells that show high pERK levels regardless of serum-free starvation. Overlay of two flow cytometry 2D contour plots of cells transfected to express high amounts of BRAF-mEGFP, after two hours of starvation. Data after staining with a concentration matched rabbit IgG antibody targeting mCherry are in gray, and data after staining with rabbit IgG pERK antibody are in green.

**Figure S9.**
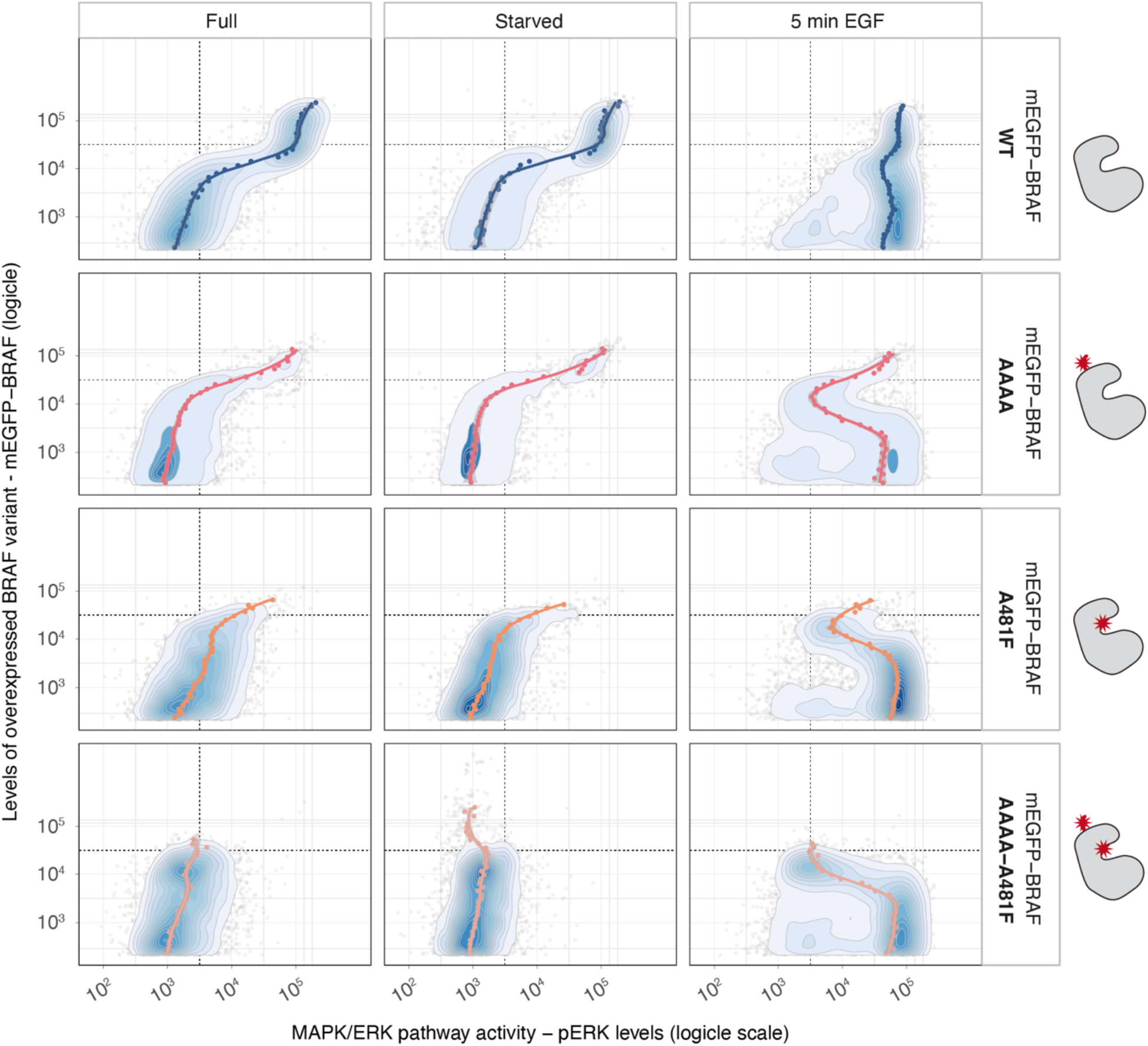
Dose response of MAPK/ERK activity (pERK levels) to different BRAF variants in HEK293T cells – replicate 2. Data is collected as in Figure 3 and represents a biological replicate as the cells were seeded, transfected, starved, stimulated, fixed, stained, and analyzed on a different day.

**Figure S10.**
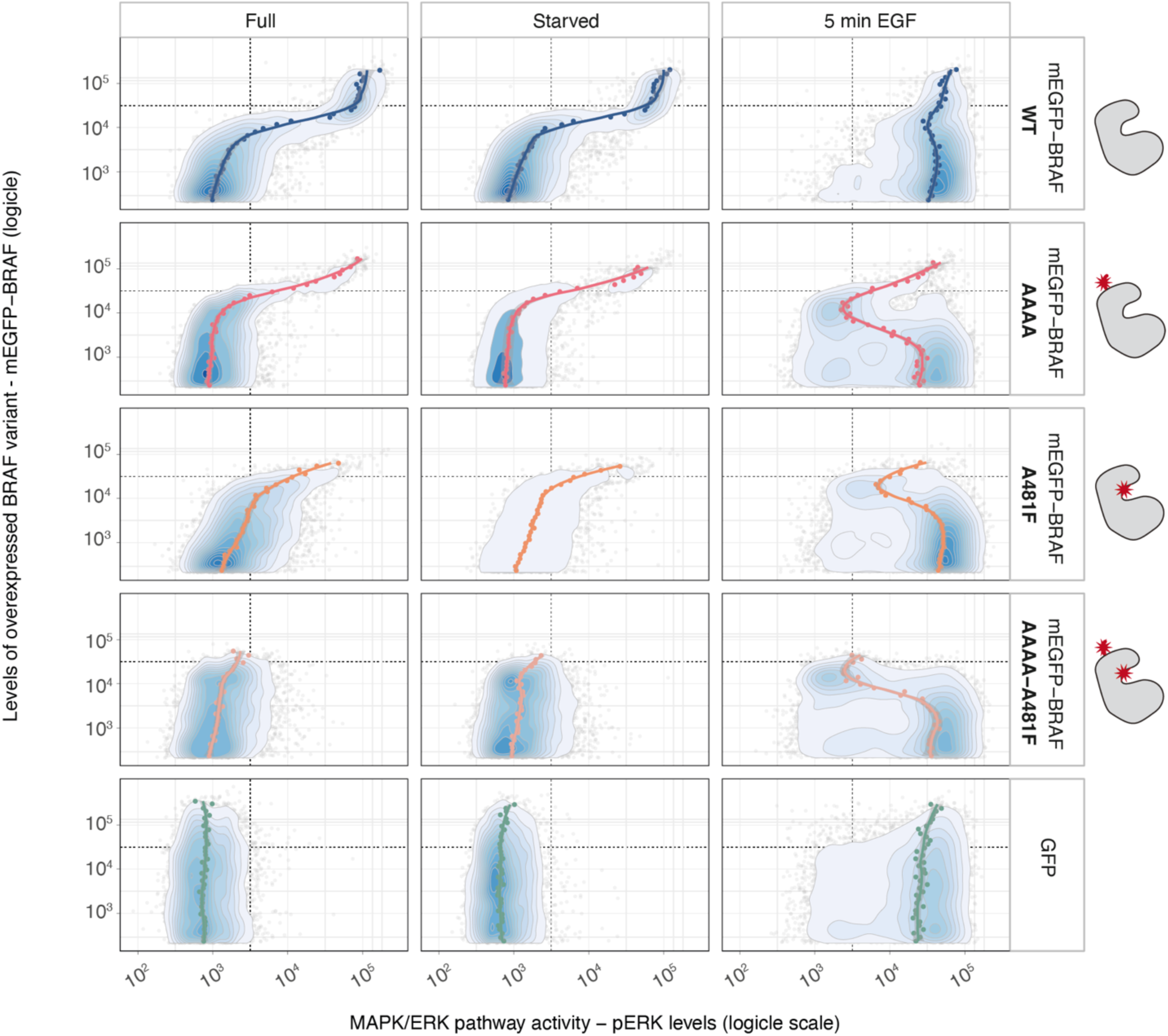
Dose response of MAPK/ERK activity (pERK levels) to different BRAF variants in HEK293T cells – replicate 3. Data is collected as in Figure 3 and represents a biological replicate as the cells were seeded, transfected, starved, stimulated, fixed, stained, and analyzed on a different day.

**Figure S11.**
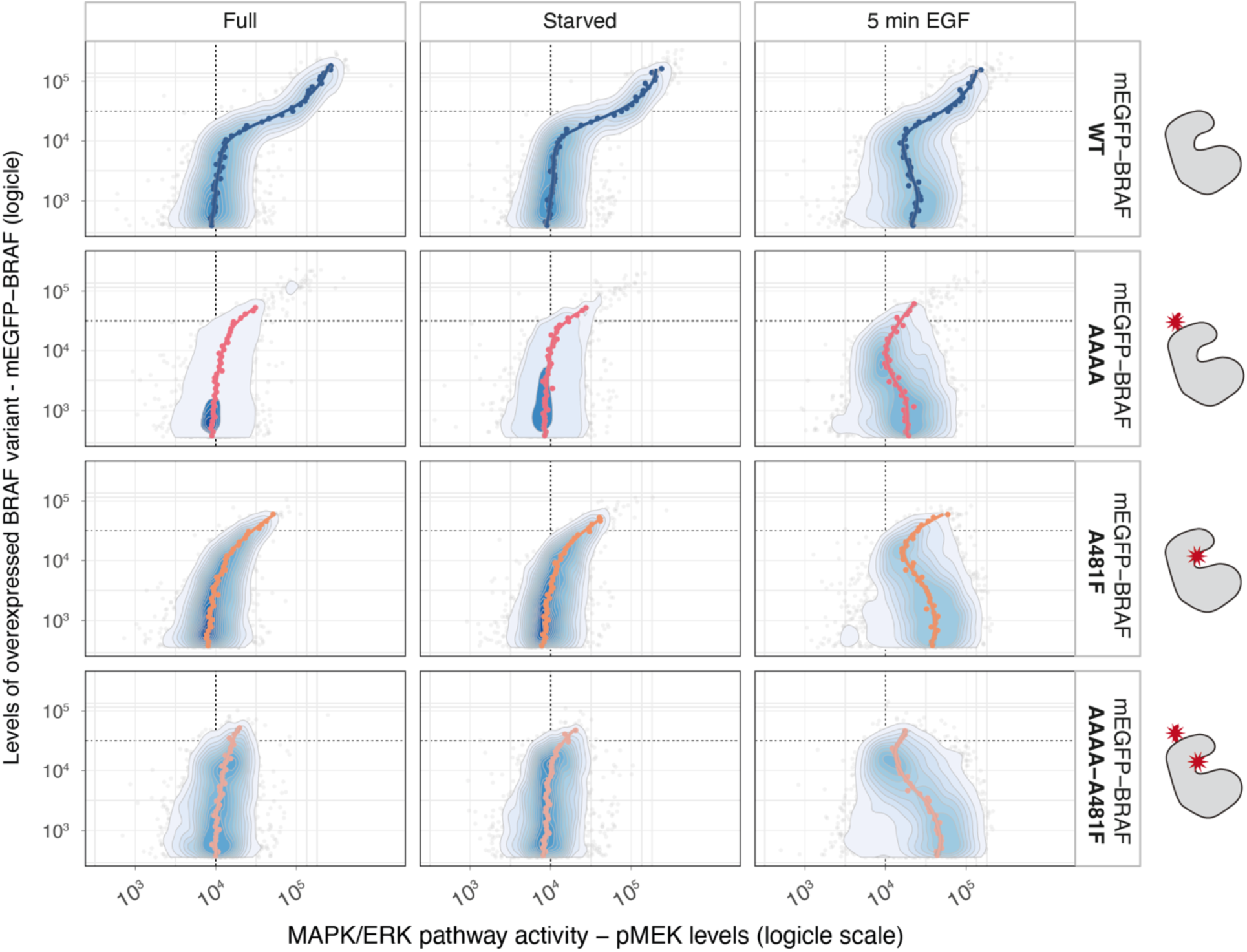
Dose response of MEK1/2 activity (pMEK levels) to different BRAF variants in HEK293T cells. pMEK levels are quantified with a phospho-MEK1/2 (Ser221) primary antibody followed by a secondary Alexa Fluor 647 conjugate antibody. Protein levels of overexpressed BRAF variants are quantified via fluorescence of the amino-terminal monomeric mEGFP tag. The experiment is otherwise performed the same as in Figure 3.

**Figure S12.**
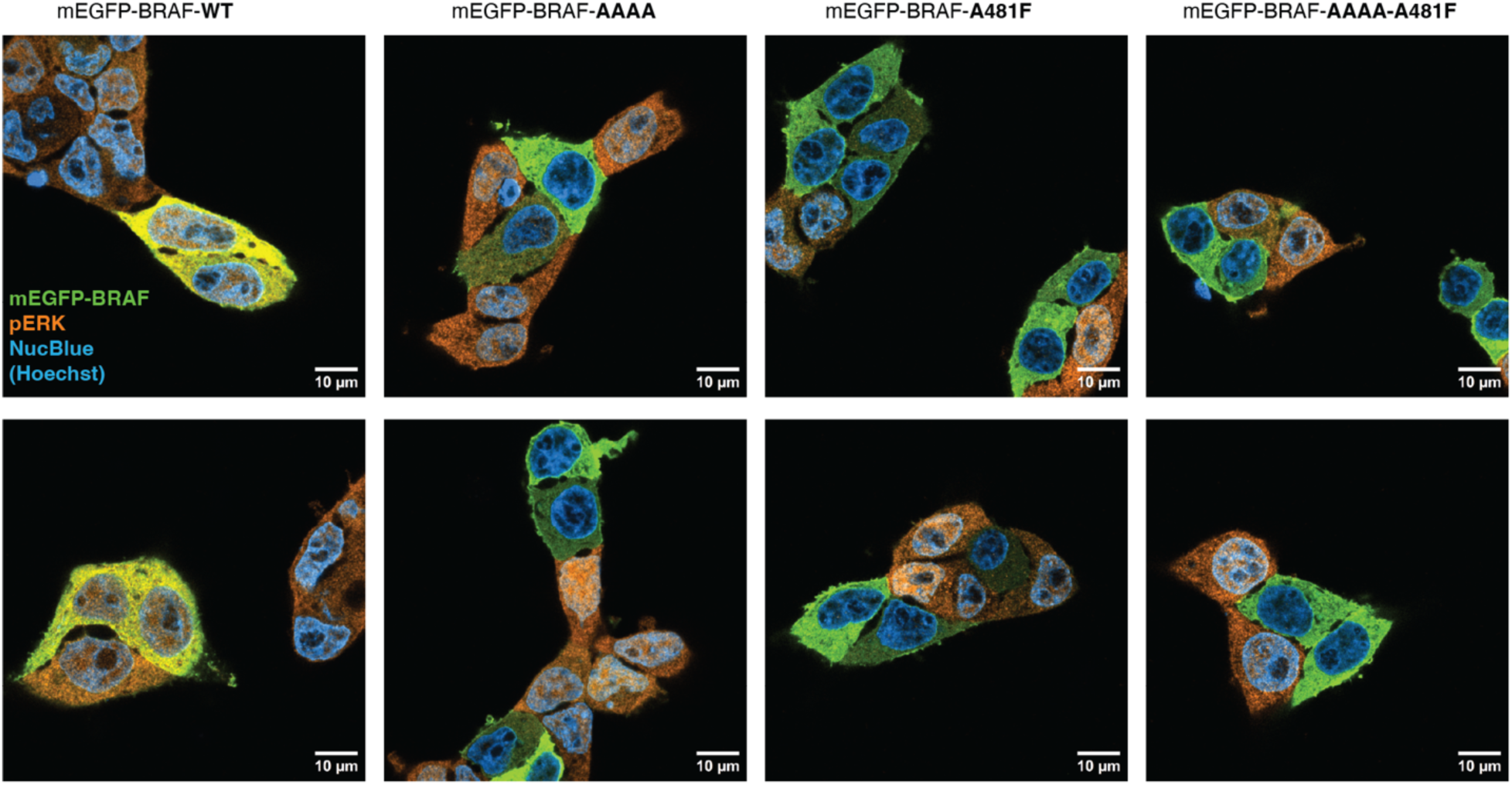
Selected confocal microscopy images of cells fixed 5 minutes after stimulation with 30 ng/ml EGF. Active MAPK/ERK is quantified with an anti-pERK antibody, and mEGFP-BRAF variant levels by GFP fluorescence. Cells were prepared and treated the same as in Figure 3 (5 min EGF condition) but imaged instead of detached for flow cytometry analysis. GFP is in green and pERK in orange. Cells that simultaneously show high levels of GFP and pERK are yellow.

**Figure S13.**
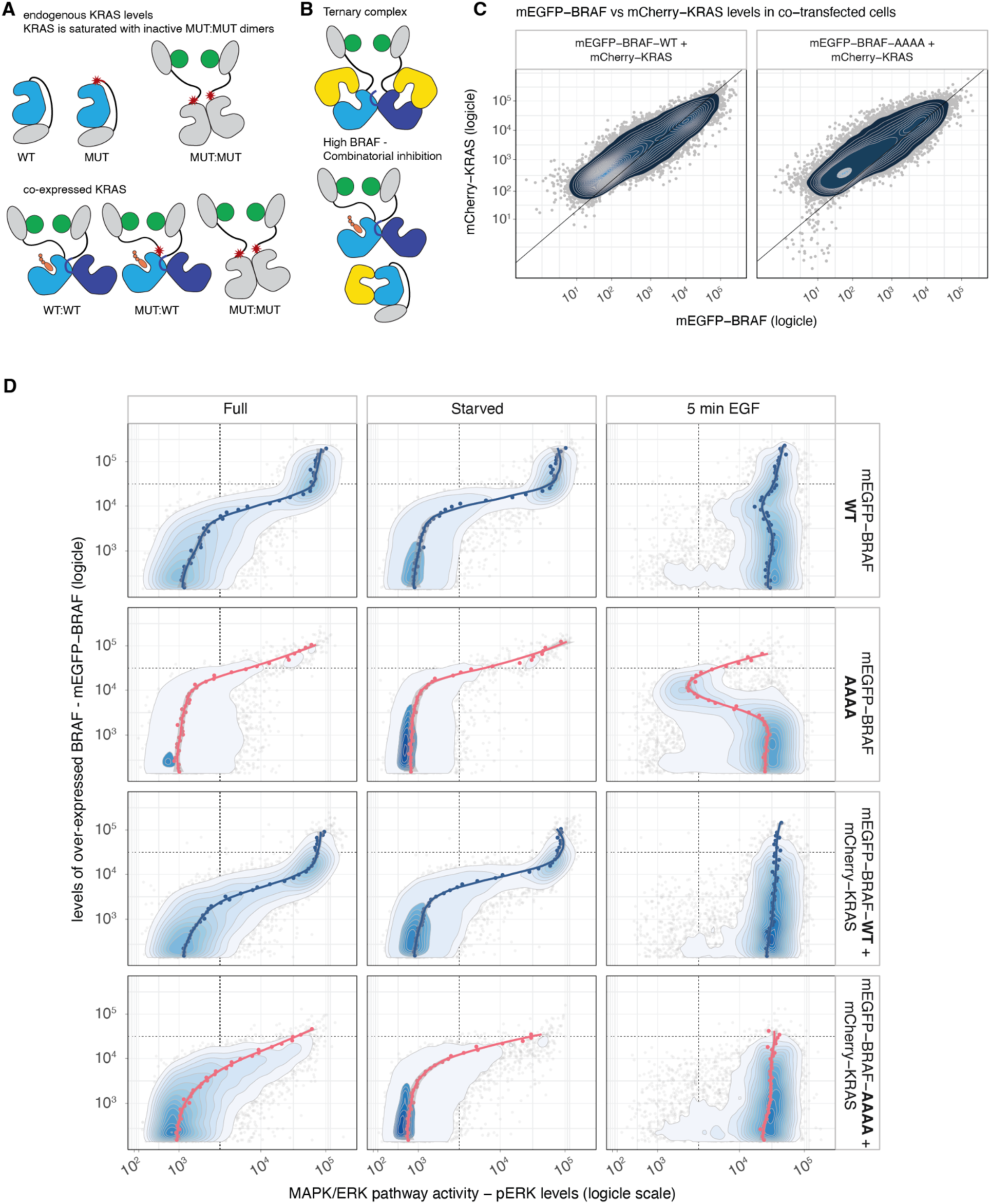
Dose response of MAPK/ERK activity (pERK levels) to levels of an activator-abrogating BRAF mutant (mEGFP-BRAF-AAAA) when KRAS is co-expressed with BRAF. Kinase schemes are as in Figure S1, with the membrane and 14-3-3 dimer omitted for simplicity. (A) When KRAS is co-expressed with BRAF-AAAA, the cellular KRAS pool does not get saturated with inactive MUT:MUT heterodimers. (B) Activating phosphorylation of ERK1/2 requires an active KRAS:BRAF:MEK ternary complex to be formed. Overexpression of BRAF while KRAS remains at endogenous levels can lead to combinatorial inhibition. (C) Expression levels of mCherry-KRAS and mEGFP-BRAF correlate in HEK293T cells co-transfected with two plasmids. (D) Flow cytometry analysis of single cells expressing mEGFP-BRAF-WT or mEGFP-BRAF-AAAA, both with or without co-transfection with mCherry-KRAS, stained for pERK. Flow cytometry data collected as in Figure 3. Values are plotted on a logicle scale. Samples gated for single cells are shown as contour plots with outlier cells represented as individual points, where blue corresponds to highest density and white to lowest density of cells. The trend lines for each condition are colored by BRAF variant (blue for wild type, pink for BRAF-AAAA) and represent the median pERK levels per each of the 50 GFP level bins of equal length. pERK activity is quantified with a pERK (pThr202/pTyr204) primary antibody followed by a secondary Alexa Fluor 647 conjugate antibody. Protein levels of overexpressed BRAF variants are quantified via fluorescence of the amino-terminal mEGFP tag.

**Figure S14.**
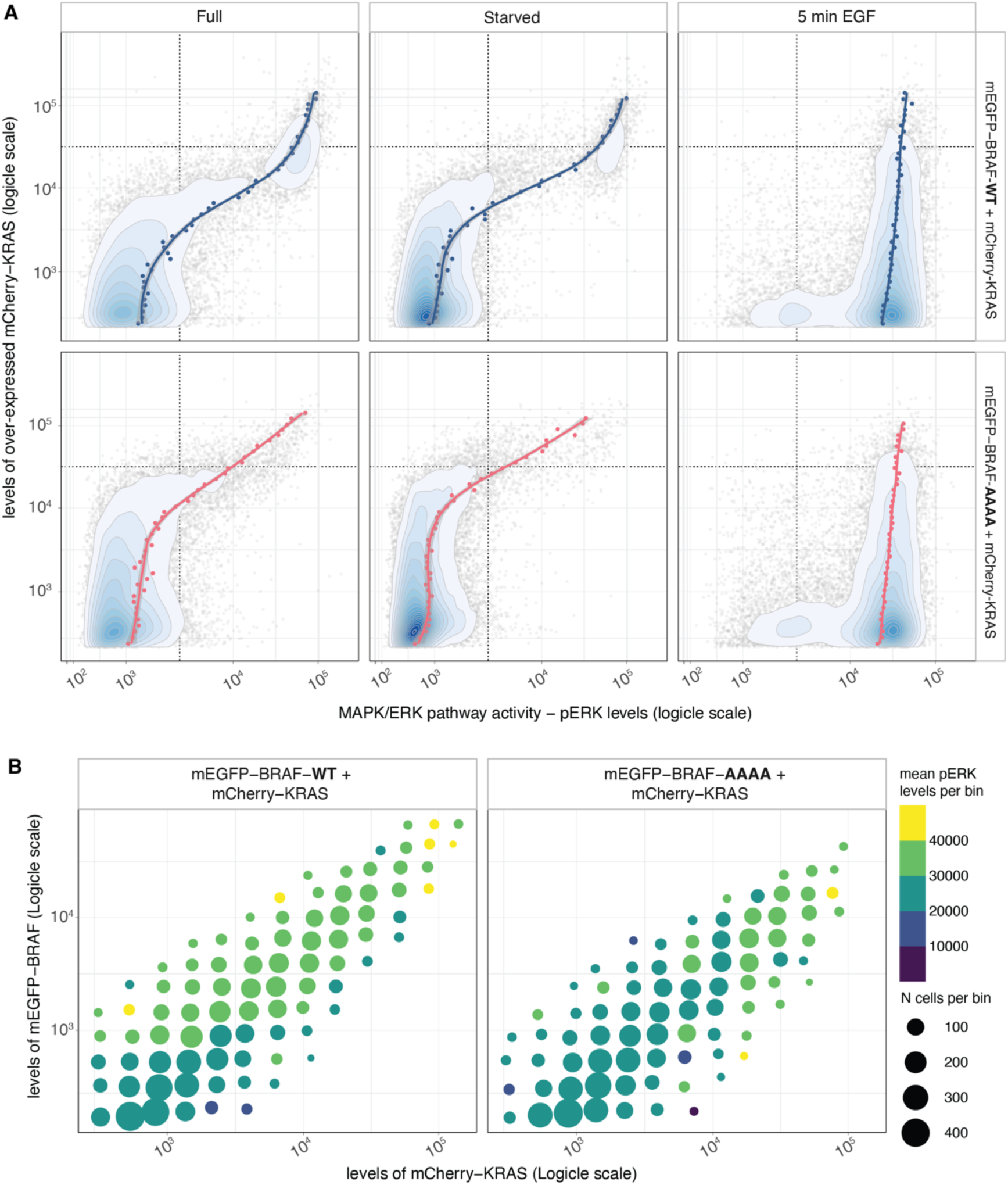
Dose response of MAPK/ERK activity (pERK levels) to levels of overexpressed mCherry-KRAS when co-transfected with mEGFP-BRAF-WT or mEGFP-BRAF-AAAA. (A) Flow cytometry analysis of single cells expressing mCherry-KRAS co-transfected with mEGFP-BRAF-WT or mEGFP-BRAF-AAAA, stained for pERK. Flow cytometry data collected as in Figure 3. Values are plotted on a logicle scale. Samples gated for single cells are shown as contour plots with outlier cells represented as individual points, where blue corresponds to highest density and white to lowest density of cells. The trend lines for each condition are colored by BRAF variant (blue for wild type, pink for BRAF-AAAA) and represent the median pERK levels per each of the 50 mCherry level bins of equal length. pERK activity is quantified with a pERK (pThr202/pTyr204) primary antibody followed by a secondary Alexa Fluor 647 conjugate antibody. Protein levels of overexpressed KRAS are quantified via fluorescence of the amino-terminal mCherry tag. (B) Flow cytometry data of cells co-transfected with mCherry-KRAS and either mEGFP-BRAF-WT or mEGFP-BRAF-AAAA and stimulated with EGF, as above and in Figure S14. The data is binned by mCherry and EGFP levels and each bin is colored by mean pERK levels.

**Figure S15.**
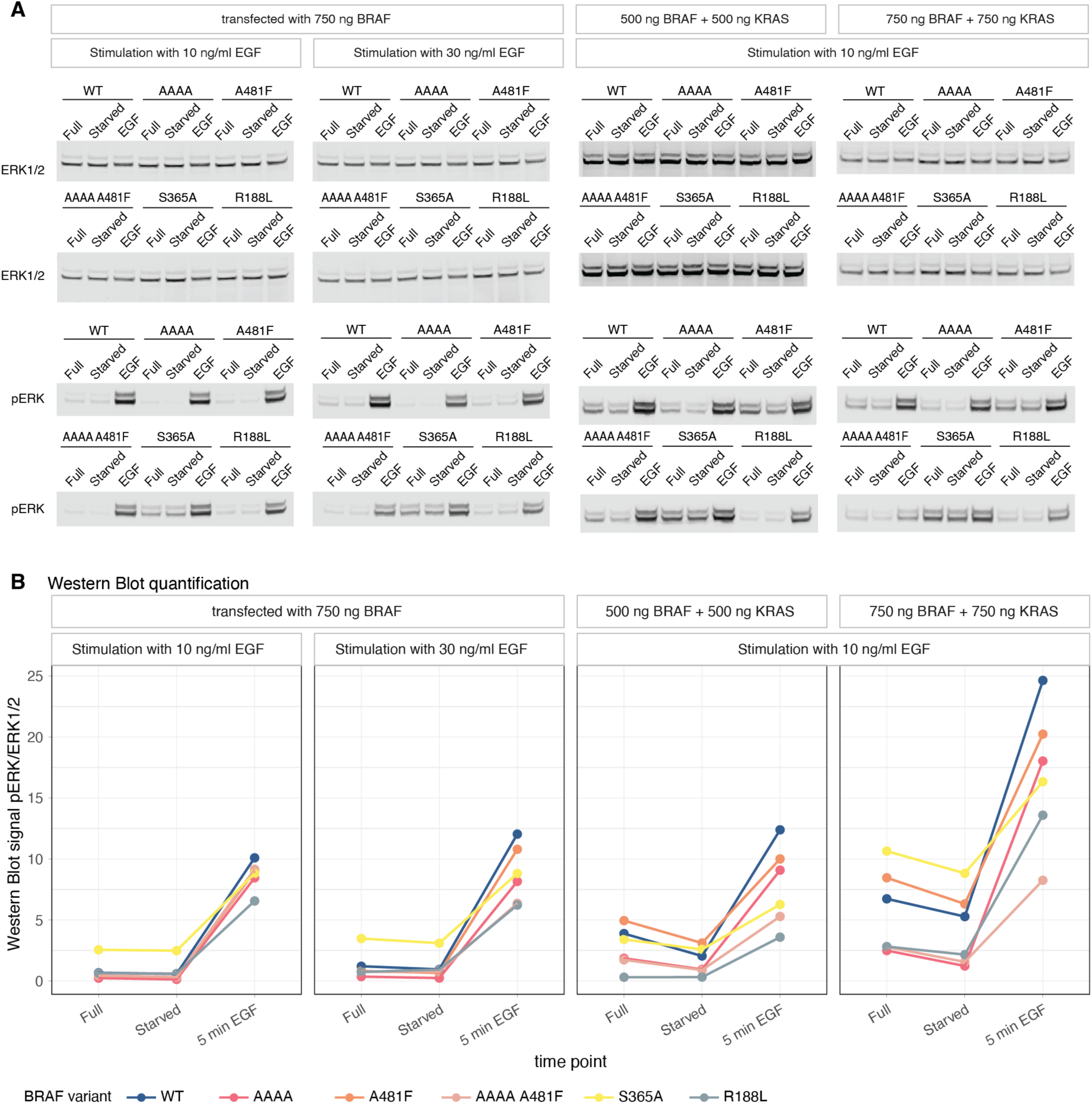
Bulk measurements by Western blot of pERK levels in cells expressing BRAF variants that abrogate activator (BRAF-AAAA), receiver (BRAF-A481F), or both (BRAF-AAAA-A481F) functions in EGF stimulated HEK293T cells. (A) HEK293T cells were transfected with 500 or 750 ng of plasmid expressing mEGFP-BRAF with or without co-expression of mCherry-KRAS and grown in medium with 10% serum (Full). After 2 hours of starvation in medium without serum (Starved), cells were stimulated with 10 or 30 ng/ml EGF (EGF). ERK1/2 and pERK levels were detected with an anti-ERK1/2 and anti-pERK antibody, respectively, from the same band to enable a quantitative comparison between samples. Quantification of Western blot signal from (A). pERK levels normalized to ERK1/2 quantified by Western blot.

**Figure S16.**
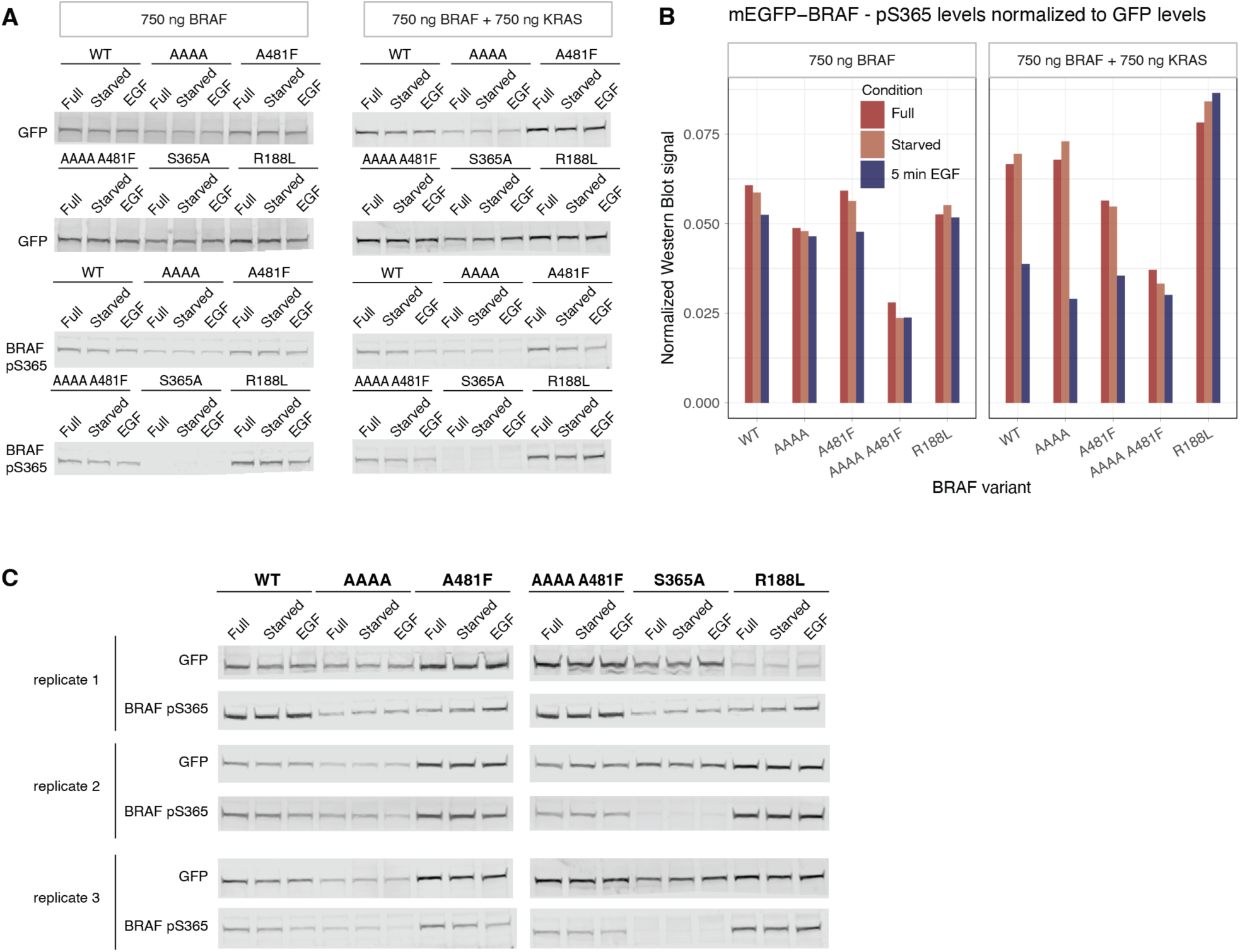
Quantification of BRAF Ser365 phosphorylation and dephosphorylation in HEK293T cells upon EGF stimulation by Western blot. (A) Co-expression of KRAS (mCherry-KRAS) is necessary to observe pSer365 dephosphorylation in overexpressed BRAF (mEGFP-BRAF) in HEK293T cells. Levels of mEGFP-BRAF variants (with anti-GFP antibody) and BRAF pSer365 were quantified in HEK293T cells grown in media with 10% FCS (Full), in cells starved for 2 hours in media without FCS (Starved), and in cells stimulated after 2 hours starvation with 10 ng/ml of EGF for 5 minutes (EGF). BRAF pSer365 was detected using the anti phospho-CRAF pSer259 antibody that can detect overexpressed pSer365 in overexpressed BRAF. GFP and pSer365 levels were detected from the same respective bands enabling quantitative comparisons. The blots for the BRAF+KRAS condition are the same as in (C) replicate 3. (B) Quantification of Western blot signal from (A). (C) Three independent biological replicates (separate HEK293T transfections and separate Western blots) of quantification of BRAF pSer365 dephosphorylation upon EGF stimulation. Cells were co-transfected with mCherry-KRAS and different mEGFP-BRAF variants. BRAF and pSer365 levels were quantified as in (A). Plot with the quantification of these blots is in Figure 4E.

**Table S1.** Data collection and refinement statistics.

|  | symmetric BRAF:MEK1<br>heterotetramer with AMPPNP | asymmetric BRAF:MEK1<br>heterotetramer with AMPPNP | asymmetric BRAF:MEK1<br>heterotetramer with ADP |
| --- | --- | --- | --- |
| **Data Collection** |  |  |  |
| Space Group | P 2<sub>1</sub> 2<sub>1</sub> 2<sub>1</sub> | P 2<sub>1</sub> 2<sub>1</sub> 2<sub>1</sub> | P 2<sub>1</sub> 2<sub>1</sub> 2<sub>1</sub> |
| Unit-cell parameters |  |  |  |
| *a*, *b*, *c* (Å) | 87.63, 163.87, 201.10 | 66.72, 67.91, 292.54 | 68.03, 68.15, 291.74 |
| $\alpha$, $\beta$, $\gamma$ (°) | 90.0, 90.0, 90.0 | 90.0, 90.0, 90.0 | 90.0, 90.0, 90.0 |
| Resolution* (Å) | 48.06 - 2.91 (3.09 - 2.91) | 146.27 - 2.60 (2.93 - 2.60) | 49.75 - 2.30 (2.48 - 2.30) |
| *R*<sub>merge</sub> | 0.080 (1.404) | 0.260 (1.663) | 0.055 (1.317) |
| Completeness (%) |  |  |  |
| spherical | 79.0 (24.4) | 56.3 (9.5) | 78.8 (20.4) |
| ellipsoidal | 94.6 (64.2) | 92.3 (73.9) | 93.3 (58.2) |
| Multiplicity | 6.8 (7.0) | 13.4 (13.3) | 6.7 (6.5) |
| *I* / σ*I* | 14.7 (1.4) | 8.1 (1.5) | 16.9 (1.4) |
| CC<sub>1/2</sub> | 1.000 (0.583) | 0.994 (0.719) | 0.999 (0.541) |
| Best/Worst diffraction<br>limits after cut-off | 2.91 / 3.78 | 2.60 / 3.79 | 2.30 / 2.98 |
| **Refinement** |  |  |  |
| Resolution (Å) | 48.06 - 2.91 (3.02 - 2.91) | 60.70 - 2.60 (2.72 - 2.60) | 49.75 - 2.30 (2.48 - 2.30) |
| No. reflections | 50612 (885) | 23610 (123) | 48275 (510) |
| *R*<sub>work</sub> / *R*<sub>free</sub> | 0.205/0.250 | 0.208/0.262 | 0.211/0.253 |
| No. atoms | 17739 | 9350 | 9218 |
| Solvent | 30 | 59 | 143 |
| Rotamer Outliers (%) | 1.72 | 2.57 | 1.92 |
| RMSD bond lengths (Å) | 0.004 | 0.002 | 0.004 |
| RMSD bond angles (°) | 0.69 | 0.55 | 0.75 |
| Average B factor (Å<sup>2</sup>) | 107.78 | 50.7 | 84.8 |
| Clash score | 8.77 | 9.62 | 9.26 |
| Ramachandran<br>favoured/outliers (%) | 96.89/0.46 | 95.98/0.09 | 96.53/0.91 |
| PDB ID | 9RTQ | 9RTR | 9RTS |

1103 \*Values of the highest resolution shell are given in parentheses

**Table S2.**
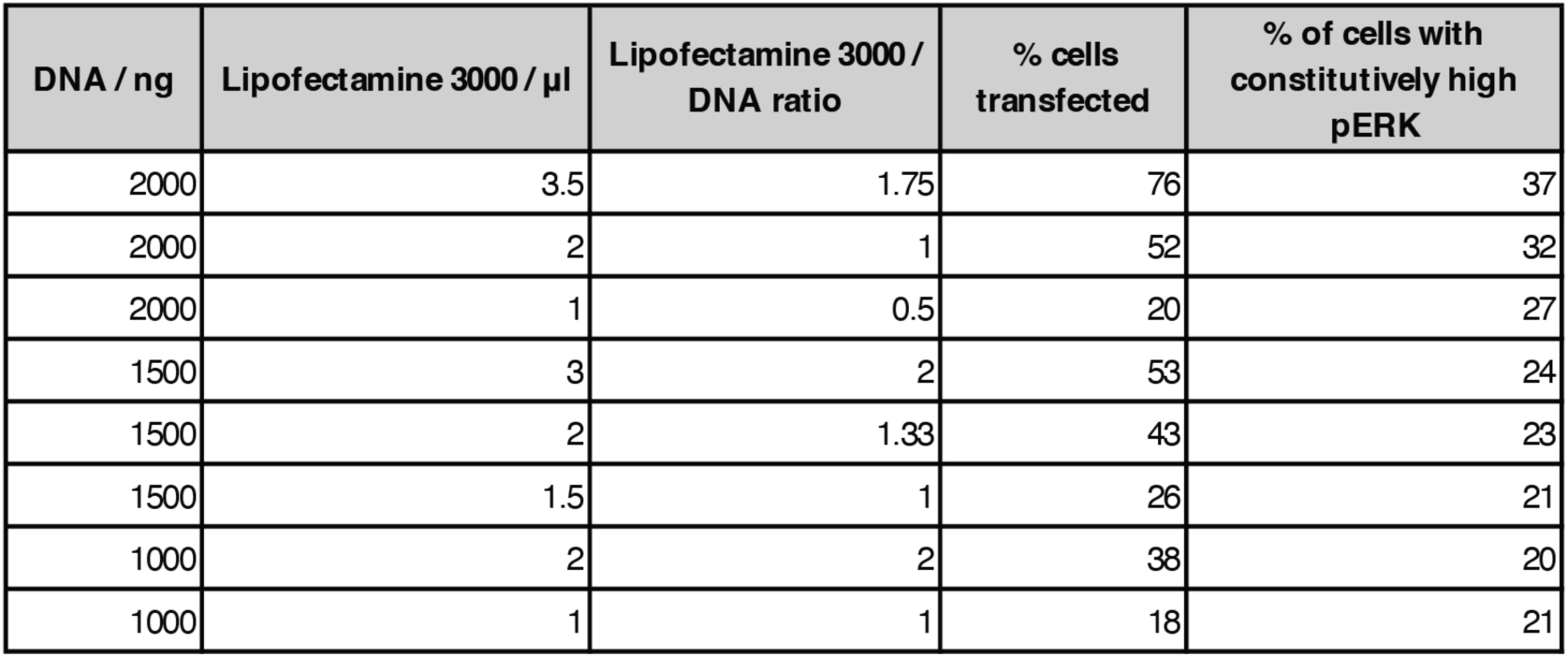
Optimization of EGFP-BRAF expression from the PGK promoter in HEK293T cells.

## METHOD DETAILS

### Protein Preparation

The BRAF kinase domain with 14 mutations was cloned into the 2O-T vector, a gift from Scott Gradia (#29710, Addgene). The coding sequence of a N-terminal 6xHis tag followed by the TEV protease site and ^Δ60^MEK1 (61-393) was synthesized and cloned into the pET15b bacterial expression vector (Genscript). The expression vector for ^Δ36^MEK1 (37-393) was generated by cloning the synthesized DNA fragment into the ^Δ60^MEK1 vector using In-Fusion Snap Assembly (#638947, Takara Bio). BRAF and MEK1 constructs were overexpressed in chemically competent Bl21(DE3) *E. coli* cells. Expression was induced by adding 0.5 mM IPTG to the bacterial culture grown in 2xYT media (Lab Logistics Group GmbH) when the optical density at 600 nm was between 0.4 and 0.6. The protein was expressed at 18°C overnight before the cells were harvested by centrifugation. The cell pellet was resuspended in Ni-A buffer (20 mM Tris pH 8.0, 500 mM NaCl, 500 mM Urea, 0.5 mM TCEP) and stored at -80°C.

The cells expressing MEK1 were lysed by sonication with VCX-600 (SONICS & MATERIALS, Inc.). Cell lysate was cleared by ultracentrifugation at 142,000g for 35 minutes. The cleared lysate was applied to a HisTrap column (GE Healthcare) equilibrated with Ni-A buffer using a MINIPULS3 (Gilson) peristaltic pump. The column was washed with Ni-A buffer supplemented with imidazole in a stepwise manner, and the protein was eluted with 400 mM imidazole. The fractions were analyzed by an SDS-PAGE gel and the fractions containing MEK1 were pooled. TEV protease was added at a 1:100 ratio. The protein solution was dialyzed against the Ni-A2 buffer (20 mM Tris pH 8.0, 250 mM NaCl, 0.5 mM TCEP) overnight. The dialyzed sample was applied to a HisTrap column equilibrated with Ni-A2 buffer. The same protocol as above was applied to wash and elute the proteins from the column. The flowthrough and wash fractions containing MEK1 were collected. The pooled fraction was concentrated by an Amicon-Ultra centrifugal concentrating unit (Millipore) using 3000g, flash-frozen in liquid nitrogen, and stored at -80°C.

The cells expressing BRAF constructs were lysed and applied to a HisTrap column as described above. When BRAF was eluted from the HisTrap column, purified MEK1 was added to the sample in a 1:1 molar ratio. After adding the TEV protease in a 1:100 ratio, the sample was dialyzed overnight against the Ni-A2 buffer. The protein complex was purified on the HisTrap column, and the flow-through and wash fractions containing BRAF:MEK1 complex were pooled. The complex was concentrated as described above then applied to a HiLoad 16/600 Superdex 200 pg size exclusion column (GE Healthcare) equilibrated with the SEC buffer (20 mM Tris pH 8.0, 150 mM NaCl, 0.5 mM TCEP). The elute was concentrated to 10 mg/mL as described above and the protein was flash-frozen in liquid nitrogen, and stored at -80°C.

### Crystallization

Nucleotides (2 mM) and magnesium chloride (10 mM) were added to the purified BRAF:MEK1 protein complex before setting up crystallization trays using Mosquito (SPT Labtech). The BRAF:MEK1 symmetric dimer crystals were obtained with a crystallization buffer containing 10% PEG4000, 0.1 mM NaMES pH 6.3, and 200 mM calcium acetate. Crystallization buffers for the BRAF:MEK1 asymmetric dimer crystals were as follows: BRAF:^Δ36^MEK1 with AMPPNP – 8% PEG20K/PEG550MME, 0.1 M Tris pH 8.6, 200 mM potassium bromide; BRAF:^Δ60^MEK1 with ADP – 6% PEG20K/PEG550MME, 0.1 M Tris pH 8.7, 100 mM calcium acetate. Crystals were cryoprotected by the reservoir solution supplemented with 25% glycerol.

### Structural determination

X-ray diffraction datasets were collected at PXI and PXII beamlines at the Swiss Light Source (Villigen, Switzerland) and I04 beamline at the Diamond Light Source (Didcot, UK). Diffraction patterns were indexed, integrated, and scaled by XDS^59^ or autoPROC^60^ and anisotropically truncated using the STARANISO server.^24^ Phases were solved by molecular replacement using Phaser.^61^ Structural models were edited using COOT,^62^ and the models were refined using phenix.refine.^63^ Dimer interface was analyzed by PDBePISA.^64^ Figures with structural models and electron density maps were prepared using PyMOL (Schrodinger LLC), and the domain rotations were measured using the “Angle between domains” PyMOL script (https://github.com/speleo3/PyMOL-psico).

### Tissue Culture

All experiments were performed in HEK293T cells (#ACC 635, DSMZ) grown at 37°C with 5% CO2, in DMEM with high glucose (4500 mg/L), sodium pyruvate and GlutaMax^TM^ (#31966-021, Gibco), and supplemented with 10% Fetal Calf Serum (FCS, #2-01F10-I, BioConcept) and 100 U/ml Penicillin and 0.1 µg/ml Streptomycin (#P4333, Sigma-Aldrich). Cells were always washed with sterile PBS (#D8537-500ML, Sigma-Aldrich) and dissociated with trypsin (#25300-054, Gibco). Cells were regularly monitored for mycoplasma infection.

The conditions described throughout the paper are either: Full (DMEM with high glucose, sodium pyruvate and GlutaMax^TM^, 10% FCS, and 100 U/ml Penicillin and 0.1 µg/ml Streptomycin); Starved (cells were starved for 2 hours at 37°C with 5% CO2 in DMEM with high glucose, sodium pyruvate and GlutaMax^TM^, but without FCS or Penicillin/Streptomycin); or 5 min EGF (EGF stimulation was performed with a given concentration of human recombinant Epidermal Growth Factor (EGF, #AF-100-15-1MG, Peprotech) in DMEM without serum and 0.1% Bovine Serum Albumin (BSA, #ACRO268131000, VWR)).

For imaging and Western blot experiments, due to several rounds of cell washing, tissue culture plates were pre-coated with 0.05 mg/ml solution of poly-D-lysine (#A38904-01, Gibco) for 30 minutes, after which the plates were washed three times with molecular biology grade sterile water and dried overnight under sterile conditions.

mEGFP-BRAF protein variants and mCherry-KRAS were expressed from a plasmid adapted from pEGFP-N1 (Clontech), where the CMV promoter was replaced by a PGK promoter, and, in the case of EGFP, a A206K mutation was introduced to decrease dimerization (mEGFP).^65^

### Live cell confocal imaging of cells expressing mEGFP-BRAF

60,000 HEK293T cells were seeded to an 8 well poly-D-lysine coated IbiTreat μ-slide (#80826, ibidi) and 18 hours later transfected with either 80 ng or 125 ng of either CMV-mEGFP-BRAF or PGK-mEGFP-BRAF (equivalent to 750 or 1200 ng of DNA in a 6-well plate). Cells were transfected using a 3:1 (μl:μg) ratio of Lipofectamine 3000 (#L3000008, Invitrogen) to DNA, according to manufacturer’s recommendations. Transfection medium was exchanged after 8 hours and cells were imaged 30 hours after transfection. Medium was changed to 300 μL of HBSS-HEPES with NucBlue LiveReady (Hoechst 33342) (#R37605, Invitrogen) at least 15 minutes before imaging. Cells were imaged with a CLSM – Leica SP8 inverse Falcon using an HC PL APO CS2 63x NA 1.4 oil objective, the 405 nm diode laser for nuclear staining, and the White light laser (WLL) for mEGFP. Imaging was performed with equipment maintained by the Center for Microscopy and Image Analysis, University of Zurich.

### Western blot quantification of pERK and BRAF-pSer365 levels

300,000 HEK293T cells were seeded per well of poly-D-lysine-coated 6-well plate and transfected after 18 hours with a mixture of 500 or 750 ng plasmid encoding mCherry-KRAS, and 500 or 750 ng plasmid encoding mEGFP-BRAF (wild type, or corresponding mutants), with 1.5 μl Lipofectamine 3000 and 1 μl of p3000 (#L3000008, Invitrogen) per 500 ng of total plasmid DNA according to manufacturer’s recommendations. Approximately 24 hours after transfection, cells were washed with warm, sterile PBS and either starved for 2 hours in 1.5 ml DMEM without FCS or further grown in full media with 10% FCS. After 2 hours, cells for full and starved conditions were washed twice in ice-cold PBS and lysed in 100 μl of PBS with 1% SDS, c*O*mplete protease (#04693124001, Roche), and PhosStop phosphatase inhibitors (#4906845001, Roche). For stimulation with EGF, 1.5 mL of 0.2% BSA-DMEM medium with 2x the desired EGF concentration was added to each well to a final concentration of 10 or 30 ng/ml. After 5 minutes of stimulation, each well was washed twice with ice-cold, sterile PBS and then lysed as above. Cells were scraped off the plate and sonicated in 2 ml Eppendorf tubes at 80-90% amplitude 3-6 times for 30 seconds in a precooled UP200St ultrasonic processor (Hielscher / HuberLab). Total protein concentration was determined with a Pierce BCA Protein Assay kit (#23227, Thermo Scientific) and 40 μg of total protein were loaded per well of an SDS-PAGE gel (#NP0321BOX, Invitrogen). Gel was blotted to a 0.2 μm nitrocellulose membrane (#12023955, BioRad). Total protein was labeled with a Revert 700 total protein stain (#402-467-0973, LI-COR) and imaged with a LI-COR Odyssey CLx Imager. Prior to antibody blotting, the nitrocellulose membrane was destained with the Destaining solution, washed in water, and subsequently blocked for 90 minutes with Casein Blocking Buffer (#B6429-500ML, Sigma Aldrich) diluted in PBS. Levels of mEGFP-BRAF were quantified by incubating with anti-GFP (mouse, monoclonal, GFP (4B10) #2955, Cell Signaling, 1:1000) antibody. Phosphorylation of BRAF Ser365 was quantified using the Phospho-c-Raf (Ser259) antibody (rabbit, polyclonal, #9421, Cell Signaling, 1:1000) primary antibody that can also detect Ser365 phosphorylation in overexpressed BRAF.^66^ Total ERK1/2 and activated (phosphorylated) ERK1/2 levels were quantified with p44/42 MAPK (Erk1/2) (3A7) (mouse, polyclonal, #9108, Cell Signaling, 1:1500) and Phospho-p44/42 MAP (Erk1/2) (Thr202/Tyr204) (rabbit, polyclonal, #9101, Cell Signaling, 1:1500) antibodies. Mouse and rabbit primary antibodies were detected with a goat anti-mouse CF680 (#20065-1, Biotium, 1:20,000) and goat anti-rabbit CF790 (#20344, Biotium, 1:20,000) secondary antibody, respectively. All primary antibodies were diluted in TBS with 5% BSA and 0.1% Tween. All secondary antibodies were diluted in TBS with 0.1% Tween. All membranes from the same experiment and with the same antibodies were scanned at the same time with a LI-COR Odyssey CLx Imager. Signal was quantified with Image Studio Lite Ver. 5.2 (LI-COR) software.

### Flow Cytometry

300,000 HEK293T cells were seeded per well of a 6-well plate and 8 hours later transfected with 1200 ng of mEGFP-BRAF plasmid and 3.6 μL Lipofectamine 3000 according to manufacturer’s instructions. 16 hours after transfection, either the medium was replaced (for the full media condition) or the cells were washed once and then incubated for 2 h in 2 ml of starvation medium at 37°C. In a 96-well V-bottom plate, two wells per condition were prepared with 65 μL of 16% fresh PFA (#15710, Electron Microscopy Sciences). For the 5 min EGF condition, 400 μL medium was removed from each well and 400 μL 5x EGF (150 ng/ml, 0.1% BSA in PBS) was added dropwise to a final concentration of 30 ng/ml. Cells were incubated for 5 min at 37°C, medium was removed, and cells were immediately vigorously resuspended in 450 μL of cold PBS to detach and obtain a single cell suspension. 200 μL of suspension was immediately added to each well of 65 μL PFA to a final concentration of 4% PFA and resuspended. Full and starve conditions were prepared in the same way but without the stimulation step. Cells were fixed for 10 minutes at room temperature, then quenched with 70 μL of 5x flow buffer (10% FCS, 10 mM EDTA in PBS). Samples were spun down at 2200g for 2 min and medium was removed by flicking the V-bottom plate. Unless stated otherwise, in all subsequent steps within the V-bottom plates, the buffer was exchanged by centrifugation at 2200g for 2 minutes followed by flicking of the plate. Cells were washed once with 1x flow buffer (2% FCS, 2 mM EDTA in PBS) and then resuspended in 25 μl of flow buffer upon which 200 μL of ice cold MeOH was added (final concentration of 90%) for permeabilization, cells were immediately resuspended and incubated on ice for 30 minutes. Samples were quenched with 70 μL of 5x flow buffer and spun down for 4 minutes at 2200g. To avoid losing cells due to high concentration of MeOH, the solution was diluted by removing 150 μL of buffer from the top of the well, and replacing it with 150 μL of flow buffer, after which the cells were again spun down for 2 minutes at 2200g. This procedure was repeated three times in total. Finally, the buffer was removed in full, and cells were washed once with flow buffer.

70 μl of Phospho-p44/42 MAPK (Erk1/2) (Thr202/Tyr204) antibody (rabbit, polyclonal, #9101, Cell Signaling, 1:150 in 0.5% BSA/PBS) or Phospho-MEK1/2 (Ser221) (166F8) (rabbit, monoclonal, #2338, Cell Signaling Technology, 1:100 in 0.5% BSA/PBS) antibody was added to each sample and resuspended. Primary antibody was incubated for 1 hour at room temperature, first quenched with 200 μL of flow buffer and then washed once with flow buffer. 70 μl of goat anti-rabbit IgG (H+L) Alexa Fluor 647 conjugate (#A-21245, Invitrogen, 1:4000 in 0.5% BSA in PBS) was added to each sample, resuspended and incubated in the dark for 30 minutes. Samples were quenched with 200 μL of flow buffer, washed once with flow buffer, and finally resuspended in 150 μL of flow buffer for flow cytometry analysis.

For the tag control experiment, a myc-tagged wild type BRAF construct was cloned by Gibson assembly. Cells were transfected, starved, fixed, permeabilized and stained with pERK following the same protocol as for mEGFP BRAF, and stained with anti-myc-tag antibody (#2276, Cell Signaling Technology, 1:500, mouse), then with Alexa Fluor 488 anti-mouse secondary antibody (#A-11001, Invitrogen, 1:1000, goat).

Samples were measured on a BD FACSymphony 5L analyser at the Cytometry Facility, University of Zurich using a 488 nm and 639 nm laser for excitation with 10000 transfection events recorded for each sample. Samples were gated for cells and single cells and the single cell population exported, parameters for these populations were used for further analysis in R.

The standard Live/Dead flow is not compatible with our time course approach as it cannot be performed post-fixation. Therefore, to control for dead cells, samples were prepared, stimulated and detached as above, washed 1x with PBS, and stained with Zombie NIR (1:200 in PBS) for 15 minutes at 37°C, quenched with flow buffer, washed once, and then fixed in 4% PFA before processing and staining as above. Importantly, these samples could only be used to estimate the number of dead cells in samples from the time course experiment, and not for pERK analysis due to increased time after stimulation.

Isotype controls were performed using a concentration-matched IgG targeting mCherry (anti-mCherry [EPR20579], #ab213511, abcam), with sample preparation as for starved or stimulated cells.

Flow cytometry was performed with equipment of the Cytometry Facility, University of Zurich.

### Imaging controls for flow cytometry samples

The same protocol was followed as for preparation of the flow cytometry samples but scaled down to an 8 well poly-D-lysine coated IbiTreat μ-slide (#80826, ibidi), without the cell detachment and resuspension step. In brief: 36,000 cells were seeded per well and 8 hours later transfected with 30 µL of the same transfection mix used for flow cytometry samples (125 ng DNA, 0.25 µL P3000, 0.375 µL Lipofectamine 3000). 16 hours after transfection, cells were washed once with starvation medium, and then starved in 200 µL of starvation medium at 37°C. After 2 hours of starvation, 40 µL of starvation medium was replaced dropwise with 40 µL of 5x EGF (150 ng/ml, 0.1% BSA in PBS) to final concentration 30 ng/ml EGF. Cells were incubated at 37°C for 5 minutes, medium was removed, and 250 µL 4% PFA in PBS was added immediately to fix the cells. Cells were incubated at room temperature for 10 minutes, washed 3x with PBS, upon which 225 µL of ice-cold MeOH was added. Cells were incubated on ice for 30 minutes, MeOH was removed, and cells were washed three times with PBS. 100 µL of primary antibody (rabbit anti-phospho-p44/42 MAPK #9101, Cell Signaling, 1:600 in 0.5% BSA/PBS) was added to each well and incubated at 4°C overnight. Cells were washed 3x with PBS, upon which 100 µL of secondary antibody (goat anti-rabbit Alexa Fluor 647, #A-21245, Invitrogen, 1:1000 in 0.5% BSA/PBS) was added to each well and stained for 30 minutes at room temperature. Cells were washed three times with PBS, then stained with 200 µL NucBlue Fixed CellReady Probes (#R37606, Invitrogen, 2 drops per ml) for 10 minutes at room temperature. Cells were again washed three times with PBS and finally imaged on a Zeiss LSM980 Airyscan using a 63x Plan-Apochromat Oil objective (NA 1.4), using the SR-4Y Airyscan detector and 405nm, 488 nm, and 639 nm lasers.

### Imaging of BRAF and KRAS co-expression and colocalization analysis

For the BRAF and KRAS co-localization experiment, proteins were expressed from a pEGFP-N1 (Clontech) plasmid encoding for mCherry-KRAS-IRES2-EGFP-BRAF under a PGK promoter. The Internal Ribosome Entry Sequence (IRES) was used the ensure sufficient levels of KRAS to localize all the overexpressed BRAF in the cell.

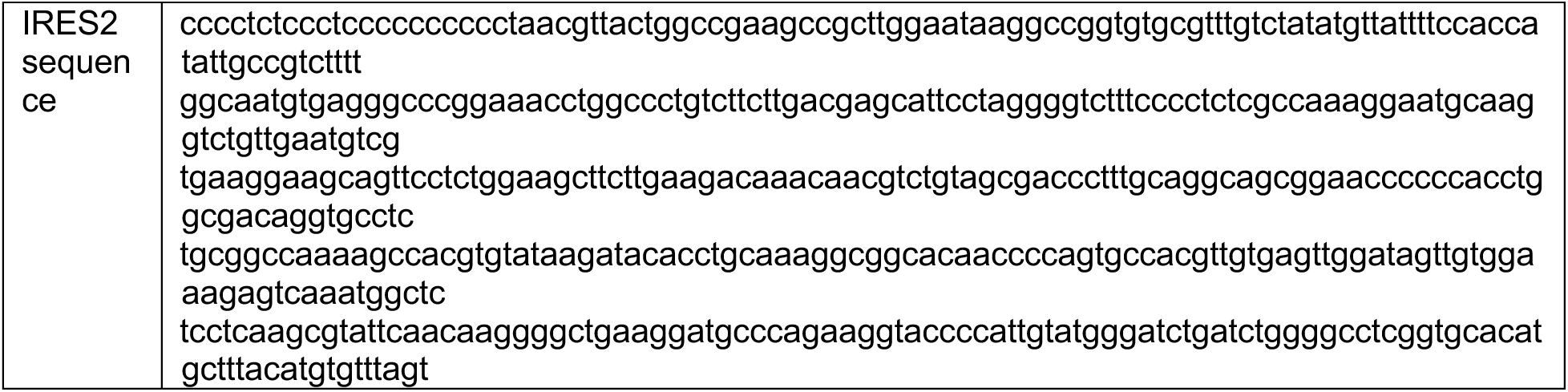
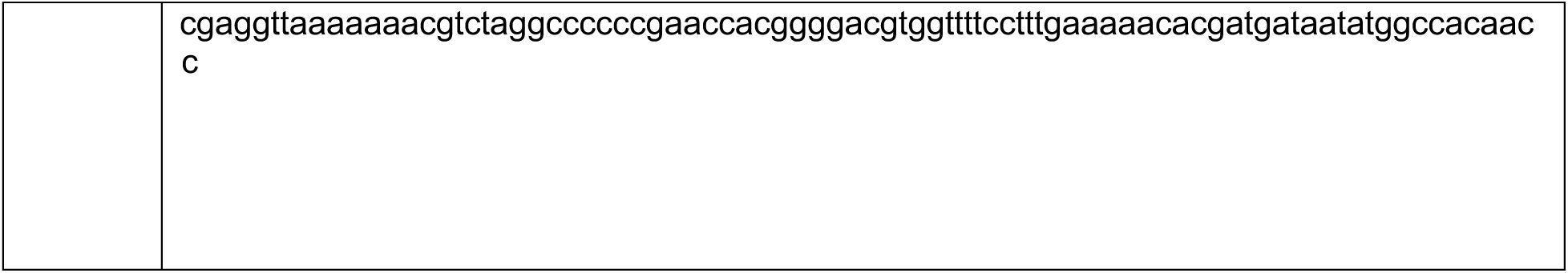

For imaging, 50,000 cells were seeded per well of a poly-D-lysine coated IbiTreat 8 well μ-slide (#80826, ibidi). After 16 hours, cells were transfected with 200 ng of mCherry-KRAS-IRES2-EGFP-BRAF plasmid and 0.5 μL Lipofectamine 3000 according to the manufacturer’s instructions. Medium was changed after 8 hours to remove the transfection mixture. 24 hours after transfection, cells were washed once with HBSS (#14175-053, Gibco) supplemented with 10 mM HEPES (#15630-056, Gibco), then stained with 200 μL 1:1000 CellMask DeepRed Plasma Membrane Stain (#C10046, Invitrogen) for 10 minutes at 37 °C. Cells were washed 3x with HBSS-HEPES, and 300 μL of HBSS-HEPES with NucBlue LiveReady (Hoechst 33342) (#R37605, Invitrogen) was added to each well at least 15 minutes before imaging. Cells were imaged on a Zeiss LSM980 Airyscan, at 37°C and 5% CO2, using a 63x Plan-Apochromat Oil objective (NA 1.4), using the SR-4Y Airyscan detector and 405nm, 488 nm, 561 nm, and 639 nm lasers. Imaging was performed with equipment maintained by the Center for Microscopy and Image Analysis, University of Zurich.

### Image analysis

A segmentation model was trained in Biodock^67^ based on the nuclear and membrane stains to segment cells while including all membrane signal. In a custom Python script, labels were loaded for each segmented cell per image, nucleus was subtracted for each cell to create a cytoplasm-membrane region of interest and mean intensity of each channel, label area, and Pearson Correlation of EGFP-BRAF and mCherry-KRAS (WT or G12D) signal were calculated for each cell in Python. Cells were filtered for transfection using a threshold on the mean KRAS and BRAF intensity assuming a 25% transfection efficiency in each condition. Furthermore, segmentation was controlled by removing cells with an area below 50 μm^2^ and via manual inspection of labels.

### Full length FLAG-GFP-BRAF purification from HEK293T cells

FLAG-mEGFP-BRAF was expressed from a plasmid adapted from pEGFP-N1 with CMV promoter (Clontech), where a A206K mutation was introduced to EGFP to decrease dimerization. 9x10^6^ cells were seeded per 15 cm tissue culture plate, two plates per construct for each biological replicate. 24 hours later, the cells in each plate were transfected with 40 μg DNA, 80 μl p3000, and 60 μl of Lipofectamine 3000. After 6 hours, media was replaced to remove the transfection mixture. 24 hours after transfection, cells were washed twice with ice-cold PBS and then scraped in 600 μl of FLAG-A buffer (100 mM Tris pH 8.0, 500 mM NaCl, 500 mM urea, 1 mM MgCl_2_) supplemented with c*O*mplete protease inhibitors (#04693124001, Roche) and flash frozen in liquid nitrogen.

The cells were lysed by sonication with VCX-600 (SONICS & MATERIALS, Inc.). The cell lysate was cleared by centrifugation at 20,913g for 10 minutes. The supernatant was incubated with anti-FLAG M2 Affinity Gel (A2220, Sigma-Aldrich) for 2 hours and applied to a Micro Bio-Spin column (Bio-Rad). The resin was washed with FLAG-A buffer followed by a wash with FLAG-A2 buffer (20 mM Tris pH 8.0, 150 mM NaCl, 1 mM MgCl_2_). FLAG-tagged mEGFP-BRAF protein was eluted with FLAG-A2 buffer supplemented with 150 µg/mL FLAG peptide (#F3290, Sigma-Aldrich) and c*O*mplete protease inhibitors.

### In vitro kinase assay

Purified BRAF and MEK1 were diluted in kinase buffer (20 mM Tris pH 8.0, 100 mM NaCl, 2 mM MgCl2, 1 mM TCEP). Final concentrations of BRAF and MEK1 were 100 nM and 5 μM, respectively. Quench buffer was prepared by mixing 4x LDS loading buffer (Invitrogen) supplemented with 10% β-mercaptoethanol and 50 mM EDTA. An aliquot of the BRAF and MEK1 mixture was mixed with the quench buffer before nucleotide was added and used as a zero time starting point. The kinase reaction was initiated by the addition of the ATP nucleotide to final concentration of 1 mM. An aliquot was taken out at each time point and the phosphorylation reaction was stopped by adding the quench buffer. Levels of phosphorylated MEK1 were quantified by Western blot following a standard protocol for Trans-blot Turbo Transfer System (Bio-Rad) with anti-phospho-MEK antibody (#MA5-15016, Invitrogen, rabbit, 1:5000 dilution) and HRP conjugated anti-rabbit IgG secondary antibody (#A27036, Invitrogen, rabbit, 1:5000 dilution). ADP with 90% and 99% purity were purchased from Sigma-Aldrich (#01897) and Thermo Scientific Chemicals (#J60672), respectively.

To phosphorylate the BRAF 14-3-3 binding sites, we used porcine PKA expressed in bacteria and prepared as described before.^51^ 100 nM of BRAF produced in mammalian cells was mixed with 100 nM of PKA. Samples at different time points were collected as above. BRAF phosphorylation was detected using the anti-phospho-CRAF (pS259) (#9421, Cell Signaling Technology, rabbit, 1:3000 dilution) and anti-phospho-BRAF (pS729) (#ab124794, Abcam, rabbit, 1:6000 dilution) antibodies.

## ACKNOWLEDGMENTS

We thank J. Kuriyan for helpful discussions. We also thank K. Nguyen and R. Dutzler for critical reading of the manuscript. We thank the support from the PSI Crystallization and the MX group at the Swiss Light Source. The authors would also like to thank Diamond Light Source for beamtime (proposal mx34035), and the staff of beamlines I04 for assistance with crystal testing and data collection. The research was funded by the Ulrich Peter & Hans Rudolf Wirz-Stiftung via Swiss Cancer Research KFS-5737-02-2023 (to Y.K. and J.S.), Swiss Innovation Agency Innosuisse Grant 42711.1 IP-LS (J.S.), Swiss National Science Foundation Sinergia Grant CRSII5_213507 (J.S.), UZH Candoc Grant FK-23-043 (J.N.), and Swiss National Science Foundation Project Grants 310030_207462 (J.S.) and 320030_227566 (T.P.).

## AUTHOR CONTRIBUTIONS

Y.K., J.N., I.N.C., and T.P. conceived the study. Y.K. performed crystallography experiments. Y.K., T.M., and J.M. performed biochemical experiments. J.N., I.N.C, G.N.D, and T.P. performed and analyzed cell-based experiments. Y.K. and T.P. wrote the manuscript with input from all coauthors. J.S. and T.P. supervised the project.

